# Directed Differentiation of Human Pluripotent Stem Cells into Radial Glia and Astrocytes Bypasses Neurogenesis

**DOI:** 10.1101/2021.08.23.457423

**Authors:** Vukasin M. Jovanovic, Claire Malley, Carlos A. Tristan, Seungmi Ryu, Pei-Hsuan Chu, Elena Barnaeva, Pinar Ormanoglu, Jennifer Colon Mercado, Sam Michael, Michael E. Ward, Anton Simeonov, Ilyas Singeç

## Abstract

Derivation of astrocytes from human pluripotent stem cells (hPSCs) is inefficient and cumbersome, impeding their use in biomedical research. Here, we developed a highly efficient chemically defined astrocyte differentiation strategy that overcomes current limitations. This approach largely bypasses neurogenesis, which otherwise precedes astrogliogenesis during brain development and *in vitro* experiments. hPSCs were first differentiated into radial glial cells (RGCs) exhibiting *in vivo*-like radial glia signatures. Activation of NOTCH and JAK/STAT pathways in *bona fide* RGCs resulted in direct astrogliogenesis confirmed by expression of various glial markers (NFIA, NFIB, SOX9, CD44, S100B, GFAP). Transcriptomic and genome-wide epigenetic analyses confirmed RGC-to-astrocyte differentiation and absence of neurogenesis. The morphological and functional identity of hPSC-derived astrocytes was confirmed by using an array of methods (e.g. electron microscopy, calcium imaging, co-culture with neurons, grafting into mouse brains). Lastly, the scalable protocol was adapted to a robotic platform and used to model Alexander disease. In conclusion, our findings uncover remarkable plasticity in neural lineage progression that can be exploited to manufacture large numbers of human hPSC-derived astrocytes for drug development and regenerative medicine.

## Introduction

Neuroglia was first described by Rudolf Virchow in 1846 as a connective substance or ‘brain glue’ that appeared to surround the nervous elements. Ramón y Cajal predicted the importance of astrocytes and dedicated a significant body of work to characterize them morphologically^1^. Today we know that astrocytes are the most abundant cell type in the central nervous system (CNS) and dynamically control a broad range of structural and functional properties of the human brain (e.g. formation of the blood-brain barrier, metabolic and trophic support, synaptogenesis and synaptic pruning, neurotransmitter uptake, ionic milieu control in the extracellular space, innate and adaptive immunity, neural repair, and others)^2-5^. Impairment of these functions are important causative or contributing factors to various neurological and psychiatric disorders including autism, epilepsy, Alzheimer’s disease, amyotrophic lateral sclerosis and others^6-8^.

During CNS development, early neuroepithelial cells give rise to RGCs, which generate large number of neurons in the cortical plate and provide a glial scaffold for migratory neurons (neurogenic phase)^9,10^. Subsequently, RGCs that remain proliferative throughout the progressive waves of neurogenesis, differentiate into astrocytes and then later into oligodendrocytes (gliogenic phase)^3,11-13^. The invariance of this developmental sequence, neurogenesis followed by gliogenesis, is highly conserved throughout vertebrate phylogeny^11^. However, despite improved understanding of how spatiotemporally orchestrated gene regulatory networks determine CNS morphogenesis, little is known about the precise cell-intrinsic mechanisms and extracellular signals that coordinate the genesis of human RGCs and their differentiation into astrocytes.

Previous reports generated astrocytes from hPSCs by prolonged culture for up to 20 months, which seemed necessary to recapitulate the neurogenic-to-gliogenic fate switch^14-16^. These strategies often required undefined media components such as fetal bovine serum (FBS) and antibody-based cell sorting approaches to enrich for astrocyte-like cells^15,17^. Moreover, to accelerate the onset of astrocyte differentiation, recent reports ectopically expressed specific transcription factors in fibroblasts and iPSC-derived NSCs^18,19^. To address these challenges, we focused on developing a method that follows the principles of developmental biology, which is stepwise and controlled differentiation, but is also scalable and efficient as required for regenerative medicine applications. We first defined conditions for neural conversion of hPSCs into *bona fide* RGCs. These pure cultures of RGCs were then coaxed exclusively into astrocytes without going through an obligated neurogenic phase. Using this efficient strategy and a robotic cell culture system, we achieved manufacturing of cryopreservable astrocytes at industrial scale.

### Efficient conversion of hPSCs into RGCs

All experiments described here were performed using human embryonic stem cell (hESC) and iPSC lines that were continuously cultured under feeder-free chemically defined conditions using E8 medium and vitronectin as a coating substrate. To standardize differentiation and avoid technical variability, hPSCs were detached using EDTA, counted, and a defined number of cells (10,000 cells/cm^2^) was plated on vitronectin-coated dishes in E8 medium supplemented with the CEPT small molecule cocktail for 24 h, which promotes optimal cytoprotection and cell survival^20^. The following day, medium was switched to Astro-1 medium (**Fig. 1a** and Methods), formulated to modulate several cell signaling pathways with importance for RGC differentiation including NOTCH and JAK/STAT^21^. After daily media changes, we noticed that differentiating cells changed their morphology, acquired more elongated cell bodies, and formed neural rosettes (day 7) expressing the RGC markers FABP7 (also known as brain lipid-binding protein, BLBP) and PAX6 (**Fig. 1b**). To confirm reproducibility of our approach, several hESC and iPSC lines were tested and showed highly efficient differentiation into RGCs expressing FABP7 and PAX6 (**Extended Data Fig. 1a and 1b**). Next, we performed RNA-seq experiments to compare our Astro-1 strategy to a widely used neural induction method, known as dual-SMAD inhibition (dSMADi), which is based on simultaneous inhibition of BMP and TGF-beta pathways^22^. This comparison revealed that neural cells generated by Astro-1 versus dSMADi substantially differed in expression of RGC genes (**Fig. 1c**). After normalization to basal expression levels in iPSCs, a total of 44 genes was found to be differentially expressed between both conditions such that 37 genes were induced by Astro-1 and 7 genes by dSMADi treatment. Notably, out of the 37 genes specifically upregulated by Astro-1, the following 15 genes were previously reported to be specifically enriched in RGCs *in vivo*^*23*^: *BNIP3, HIST1HD, HMMR, KIF20A, TRIM24, ASPM, BIRC5, CACHD1, FAT1, HIST1H1B, HIST1H1C, HIST1H1E, HIST2H2BE, LDHA, NUSAP1*. In contrast, only one RGC gene, *PSAP*, was differentially upregulated by dSMADi. FAT1 regulates the proliferation of RGCs and its loss leads to disrupted radial glia cytoarchitecture and neural tube defects^24^. By using immunocytochemistry and Western blot analysis, we found that atypical cadherin FAT1 was strongly expressed by FABP7^+^ neural rosettes **(Extended Data Fig. 2e)**. Notably, FABP7 and FAT1 proteins were expressed at 2-and 4.8-fold higher levels in cells differentiated with Astro-1 versus dSMADi, whereas PAX6 expression was at comparable levels (**Fig. 1d**, numerical values represent normalization to GAPDH). Other genes of interest are *BIRC5* (*SURVIVIN*) and *ASPM*, which control RGC function such as symmetric cell division^25,26^. Using quantitative *in situ* hybridization (RNA scope), we found that *BIRC5* transcripts were significantly higher in Astro-1 versus dSMADi (**Extended data Fig. 2a** and **2b**, *\*p=0*.*0115*). Similarly, mitotic spindle assembly protein ASPM was more abundant in cultures treated with Astro-1 versus dSMADi, although statistical analysis revealed only a trend increase (**Extended data Fig. 2c** and **2d**, *p = 0*.*24*). Collectively, these findings suggest that Astro-1 treatment differentiates hPSCs directly into RGCs.

**Fig. 1:**
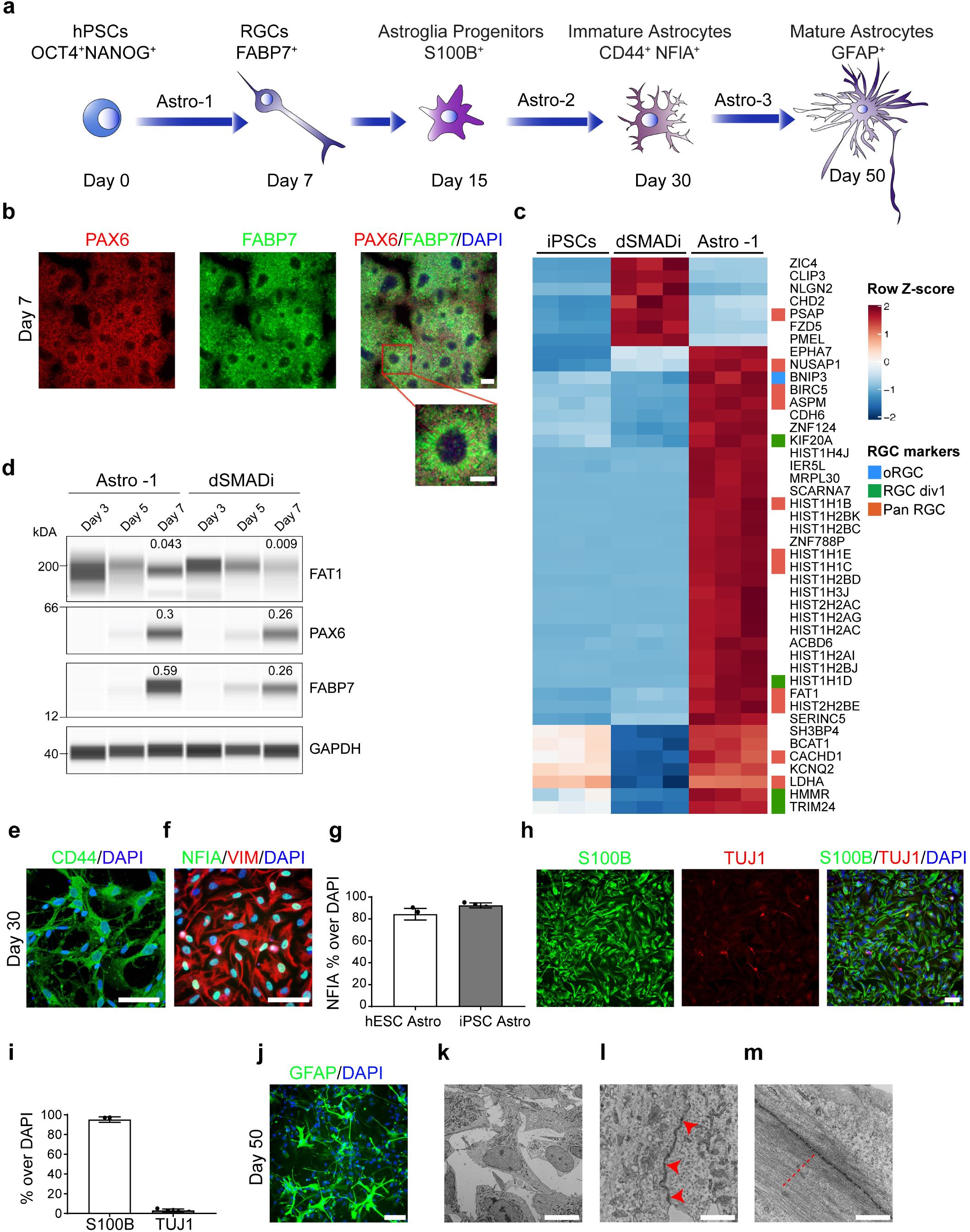
Controlled RGC and astrocyte differentiation from human pluripotent stem cells. **a**, Schematic illustration depicting RGC and astrocyte differentiation from hPSC using chemically defined Astro-1, Astro-2, and Astro-3 media (see Methods for details). **b**, Representative immunostainings of RGC cultures (day 7) showing neural rosettes expressing FABP7 and PAX6. **c**, Heatmap showing differentially expressed RGC genes after neural induction with Astro-1 and dSMADi. **d**, Quantitative Western blot analysis of FABP7 and FAT1. See the difference between cultured differentiated with Astro-1 versus dSMADi. Numerical values represent peak size normalization to GAPDH expression. **e**, Expression of CD44 by astrocytes (day 30). **f**, NFIA and VIMENTIN co-expression at day 30. **g**, Quantification of NFIA^+^ nuclei in two hPSC lines normalized to total number of nuclei at day 30. **j**, Micrograph showing astrocytes with stellate morphology expressing GFAP (day 50). **k**, Electron microscopy at day 50 confirms typical astrocyte morphologies. **l**, Depiction of adherens junctions at the ultrastructural level marked by red arrowheads. **m**, Presence of abundant intermediate filaments marked by red dashed line. Scale bars, 100 µm (**b**,**e**,**f**,**h**,**j**); 2 µm (**k**), 500 nm (**l**,**m**).

### Differentiation of RGCs into astrocytes at the expense of neurogenesis

To properly culture proliferative RGCs in Astro-1 medium, it was necessary to passage cells at defined time points to avoid overcrowded cultures that otherwise undergo cell contact inhibition and compromise directed differentiation. Hence, cells were passaged at day 7, 11, and 14 and plated at 20.000, 30.000 and 30.000 cell/cm^2^ to compensate for decreased proliferation rate over time. Cells were passaged in the presence of the CEPT cocktail, which was applied for 24 h at each passage to ensure optimal viability^20^. By day 15, nearly all cells had adopted a flat morphology and around 50% of total cells already expressed the glial marker S100B (**Ext. Data Fig. 2f and 2g**). At this stage, we removed LDN-193189 and PDGF-AA and included a chemically-defined lipid supplement to provide a more enriched medium (Astro-2 medium), while avoiding animal serum (**Fig. 1a** and Methods). Under these conditions, S100B^+^ cells were less proliferative, and passaging was carried out twice (day 18 and 23). By day 30, virtually all cells expressed the surface protein CD44 (>95%) and more than 85% of total cells expressed NFIA (**Fig. 1e-g**), an important transcription factor controlling astrogliogenesis^11,27^. At this stage, we noted a diffuse immunostaining pattern for glial fibrillary acidic protein (GFAP) (**Extended Data Fig. 2h**) but upon further maturation in Astro-3 (**Fig. 1a** and Methods), typical cell morphologies and GFAP expression was obtained (**Fig. 1j, Extended Data Fig. 1i**). Again, this observation was confirmed in multiple hESC and iPSC lines (**Extended data Fig 1c** and **1d**). To validate the absence of neurons and the purity of astrocyte cultures, we performed immunostainings and quantitative microscopic analysis revealing that more than 90% of total cells expressed the glial marker S100B by day 30, with only sporadic occurrence of TUJ1^+^ neurons (3% of total cells) (**Fig. 1h** and **1i**). Likewise, electron microscopy of hPSC-derived astrocytes at day 30 and 50 and comparison to human iPSC-derived astrocytes from a commercial vendor (FUJIFILM CDI) did not detect cells with neuronal characteristics (**Extended data Fig. 3a**). Instead, ultrastructural analysis showed star-shaped morphologies (**Fig. 1k**) and abundant intermediate filaments in astrocytic processes (**Fig. 1m**, red dashed line and **Extended Data Fig. 3c**, red lines). Cells were rich in organelles indicating secretory activity (endoplasmic reticulum, Golgi apparatus) (**Ext. Data Fig 3a and 3b**). Well-developed cell junctions such as *zonula adherens* were also confirmed in all samples (**Fig. 1l**, red arrowheads and **Extended Data Fig. 3b**, red arrowheads).

Next, we performed time-course RNA-seq experiments to characterize human astrogliogenesis under these defined conditions. Principal component analysis (PCA) showed distinct clustering of samples across different timepoints (day 0, 7, 14, 21, 30, 50) (**Fig. 2a**). Unbiased comparison of the top 100 differentially upregulated genes using the *ARCHS4* tissue database and the gene enrichment platform ENRICHR^28^, confirmed stepwise differentiation resulting in “*astrocyte*” as the top category by day 21, 30 and 50 (**Fig. 2c**). Gene expression heatmap analysis showed cell type and stage-specific gene expression and the observed transcriptional changes were consistent with stepwise astrogliogenesis (**Fig. 2b**). Accordingly, pluripotency-associated genes *OCT4* and *NANOG* were downregulated by day 7, whereas *NOTCH1, FABP7, PAX6, NESTIN (NES), NRCAM, NCAN* were strongly induced in RGCs (**Fig. 2b**) consistent with immunocytochemistry and Western blot data (**Fig. 1b-e and Extended Data Fig. 1c and 1d**). By day 14 and 21, additional radial glia markers *VIMENTIN (VIM)* and *TENASCIN C* (*TNC)* were expressed by PAX6^*+*^ RGCs. By day 30, cells expressed canonical astrocyte markers (e.g., *NFIA, NFIB, S100B, CD44*, glutamate transporters *SLC1A2/EAAT2, SLC1A3/GLAST*). Importantly, throughout the entire glial differentiation process we did not detect any significant induction of pro-neuronal transcription factors (*NEUROGENIN 1, 2 and 3, ASCL1* or *EOMES)*. In contrast, pro-astroglial transcription factors *SOX9, NFIA, NFIB*^19,29^ were strongly expressed from day 14 onward (**Fig. 2d**). Lastly, we also noted the absence of oligodendrocyte precursor markers *OLIG1* or *OLIG2* throughout the differentiation process (**Fig. 2d**). Together, these findings document that RGCs can be directly differentiated into astrocytes without going through a neurogenic phase or artificially inducing the gliogenic switch by forced expression of *NFIA* in expanded NSCs.

**Fig. 2:**
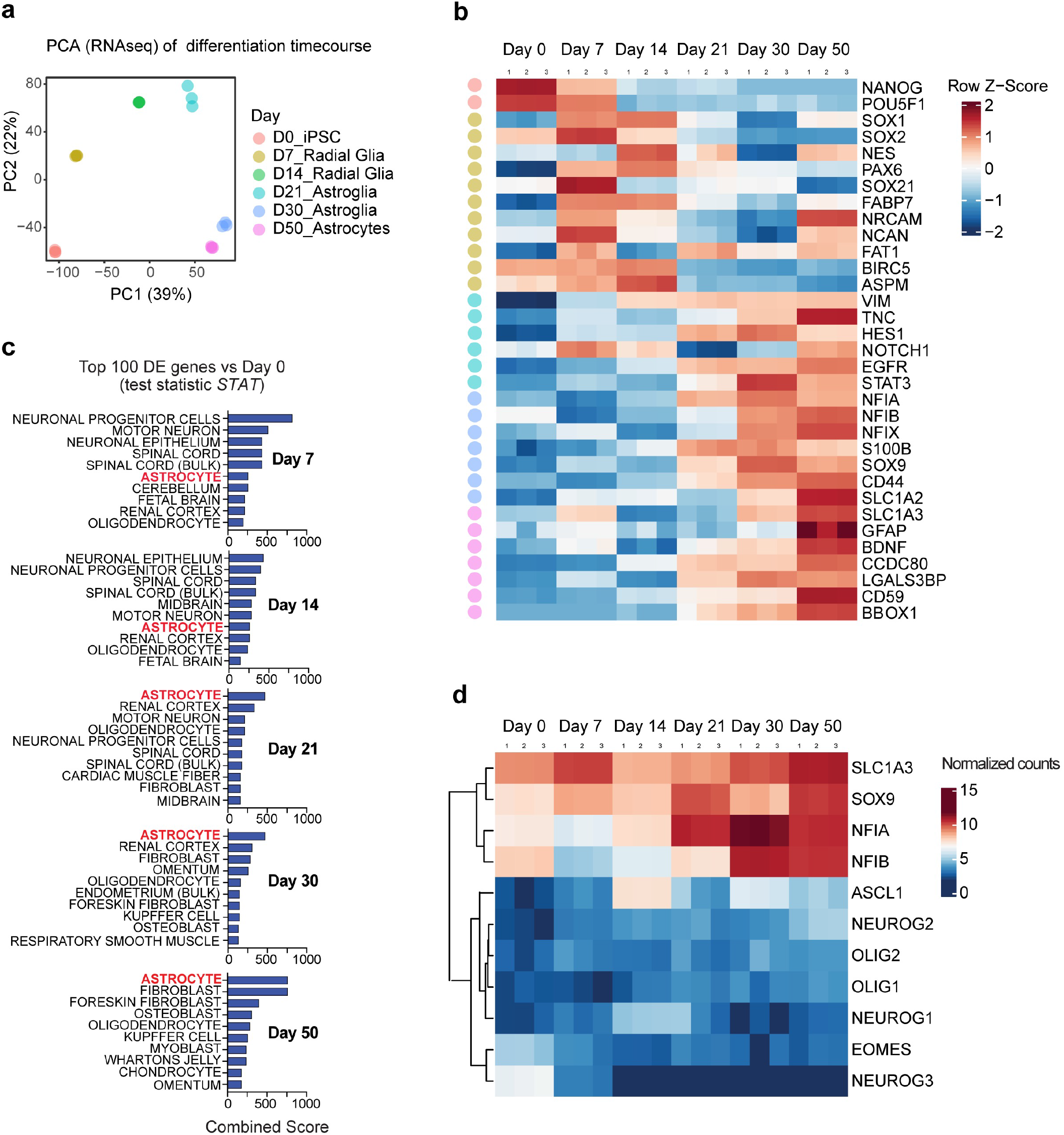
Transcriptomic analysis of hPSCs differentiating into RGCs and astrocytes. **a**, PCA of all samples used for time-course RNA-seq experiments. **b**, Heat-map depicting cell-type specific genes at different timepoints including pluripotent cells (day 0), RGCs (day 7-14), astroglia (day 21-30) and astrocytes (day 50) **c**, Gene enrichment analysis (combined score-*Enrichr*/*ARCHS4 Tissue database*) based on top 100 differentially expressed genes (*STAT value*) versus hPSCs (day 0) reveals “astrocytes” as top hit starting at day 21. **d**, Heatmap with average linkage dendrogram demonstrates predominant induction of astrogliogenic transcription factors (e.g., *SOX9, NFIA, NFIB*) and absence of genes representative for neurogenesis (*NEUROG 1/2/3, EOMES*) and oligodendrogenesis (*OLIG1/2*).

Next, we compared the transcriptomic profile of astrocytes generated with the present method to previously published reports describing hPSC-derived astrocytes^17,27,30^ and primary astrocytes from fetal and adult human brains^31^ (**Extended Data Fig. 4a**). After using the SRA batch correction method considering the expression of 50 “housekeeping” genes, we performed unsupervised hierarchical clustering based on the expression of a previously reported set of genes upregulated in human fetal astrocytes^31^ (**Extended Data Fig. 4b)**. We observed that our serum-free astrocyte cultures and hPSC-astrocytes samples from other studies clustered together with samples of primary fetal astrocytes and away from the primary adult astrocytes. Our standard serum-free cultures were exposed to a commercially available lipid supplement that is part of Astro-2 medium (see Methods), which we found to be sufficient and necessary to culture human astrocytes in the absence of animal serum. In addition, we exposed our astrocyte cultures to 2% FBS, which is typically used in other studies^17,27,30^. Interestingly, administration of 2% FBS to our astrocyte cultures led to strong induction of genes encoding histone proteins (**Extended Data Fig. 4b**-boxed in red and **Ext. Data Fig.4**), thereby clustering with primary human fetal astrocytes^31^, which was in contrast to all previously published datasets^17,27,30^.

### Epigenetic regulation of human astrogliogenesis

To characterize the epigenetic changes that occur during astrogliogenesis, we performed genome-wide analyses of chromatin accessibility (*ATACseq*) and DNA methylation (*MeDIPseq*). Starting with pluripotent cells (day 0) and harvesting samples day 7, 30 and 50, we were able to map distinct chromatin signatures as summarized in PCA plot for the ATACseq data (**Fig. 3a**). Targeted analysis for open chromatin in a set of specific RGC and astrocyte genes, showed highest chromatin accessibility for *SOX9* and *FAT1* during early differentiation (day 0 and 7), while *SLC1A2* and *SPARC* showed highest accessibility in astrocytes at day 30. *S100B* and *GFAP* were most accessible at day 50 (**Fig. 3c**). Hence, open chromatin of the *GFAP* promoter was consistent with the onset of mRNA and protein expression (**Figs. 1i, 2b** and **Extended Data Fig. 1f**). To provide further evidence for glia-restricted cell fate, we studied gene loci of the transcription factor *NEUROGENIN1 (NEUROG1)*, known to be sufficient to induce neurogenesis and inhibit RGC differentiation into astrocytes^32^. The chromatin configuration of both promoter region and gene body indicated that *NEUROG1* was inaccessible at day 30 and 50 (**Fig. 3d**). Similarly, the chromatin regions of other neurogenin family members *NEUROG2 and NEUROG3*, were also inaccessible at those time points (**Extended Data Fig. 6a, b)**. Of note, *NFIA and NFIB* were largely inaccessible at the RGC stage (day 7) but then accessible in astrocytes (day 30 and 50) (**Extended Data Fig. 6c, d)**. REST is a master regulator involved in blocking neuronal cell fate in non-neuronal cells by controlling epigenetic modifications ^33-35^. We noted that the *REST* promoter region remained accessible and *REST* transcripts, as confirmed by RNA-seq, were expressed throughout the entire differentiation trajectory (**Fig. 3e, f**). Altogether, the distinct chromatin states studied at different timepoints strongly support the notion that neurogenic potential was suppressed in favor of glial cell fate commitment. To further characterize the epigenetic mechanisms of human astrogliogenesis, we studied DNA methylation changes over the course of pluripotency to astrocyte specification (**Fig. 3g**). Of the top 50 most significant differentially methylated regions between iPSCs (day 0) and astrocytes (day 50), 49 were hypermethylated in astrocytes (**Fig. 3h**). Among these hypermethylated genes, we found pro-neuronal transcription factors IRX1 and *LHX5* ^36,37^ and several genes involved in neuronal function (*SHANK1, GPRIN1, JPH4*) ^38,39^ (**Fig. 3h**). Furthermore, sphingosine-1-phosphate receptor 5 (*S1PR5)*, a gene involved in oligodendrocyte development^40^, and *PHLDB1*, a gene associated with glioma susceptibility^41^, were also methylated in day-50 astrocytes (**Fig. 3h**). Altogether, the analysis of hypermethylated genes corroborated the notion that neuronal and oligodendroglial genes remain suppressed, while an astrocyte-specific gene expression program unfolds. Next, to gain deeper insights into the epigenetic modifications in human astrocytes, we prepared heatmaps summarizing the peaks of highest variance in promoter regions and gene bodies encompassing over 300 glial genes^31^. Specifically, we compared peaks of highest variance detected in iPSCs (day 0) and astrocytes (day 30 and 50). Interestingly, two clusters comprised of 31 and 79 genes, respectively, showed highly accessible regions in astrocytes (**Extended Data Fig. 5a)**. Gene enrichment analysis suggested that the smaller cluster with 31 genes annotated processes associated with ongoing differentiation such as cell migration, cell motility and dendrite development (**Extended Data Fig. 5b, c**). The larger cluster of 79 genes represented categories such as cell cycle and proliferation (**Extended Data Fig. 5d, e**), which may reflect the fetal-like signature of astrocytes derived from hPSCs and inform future approaches to enhance cell maturation. Next, we reasoned if the mapped temporal epigenetic changes could shed light on the cell signaling pathways that govern astrogliogenesis. Guided by the methylation changes indicated by the PCA of peaks with the highest variance, we focused on gene and promoter regions of JAK/STAT, NOTCH, PDGF, *mTOR*, and *NCOR2* (**Fig. 3h**). Interestingly, components of JAK/STAT signaling (*STAT1, STAT2, STAT3, STAT4, EGFR)*, NOTCH *(NOTCH1, NOTCH2, NOTCH3, HES4*) and tyrosine kinase receptor *PDGFRB* were consistently hypermethylated at the RGC stage and then hypomethylated in astrocytes (**Fig. 3h**). In contrast, *HES6*, which is an atypical HES gene and previously described as inhibitor of *HES1* and required for the activation of *NEUROGENIN* and *NEUROD* ^42^, was hypermethylated in astrocytes (**Fig. 3h**). Lastly, hypomethylation of *mTOR* and *NCOR2* genes specifically in GFAP-expressing astrocytes (day 50), but not in immature CD44^+^ astrocytes (day 30) or RGCs (day 7) (**Fig. 3h**), may indicate important roles in astrocyte maturation. Indeed, unbiased analysis of top differentially methylated regions and comparison immature (CD44^+^, day 30) and mature (GFAP^+^, day 50) astrocytes resulted in *NCOR2* hypomethylation as the most significant differential hit and other relevant genes are shown in the Manhattan plot as well (**Extended Data Fig. 5f**).

**Fig. 3:**
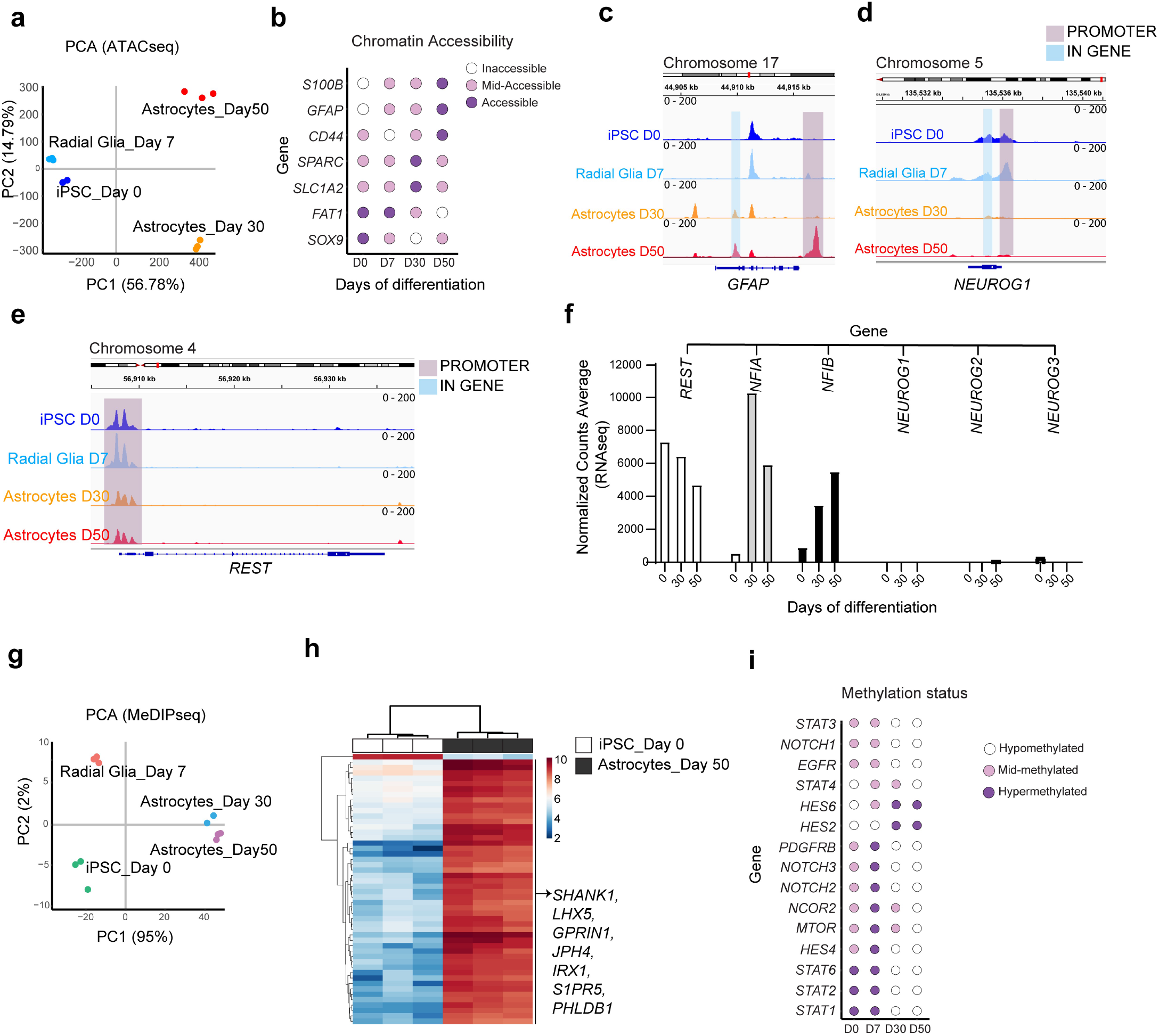
Genome-wide chromatin accessibility (ATAC-seq) and DNA methylation (MeDIP-seq) **a**, PCA plot of samples collected at different time points (day 0, 7, 30, 50) to study chromatin accessibility (ATAC-seq). **b**, Dot plot of chromatin accessibility representing peaks of highest variance in genes expressed by RGCs and astrocytes. **c**, Interactive genome viewer (IGV) plot showing peaks in *GFAP* gene loci. Note the open *GFAP* promoter region at day 50. **d**, IGV plot demonstrating changes in chromatin accessibility in *NEUROG1* gene loci. Note the global closure of *NEUROG1* region at days 30 and 50. **e**, IGV plot representing *REST* gene loci. Note that *REST* gene promoter remains accessible throughout differentiation time-course. **f**, Normalized counts (RNA-seq) of gliogenic and neurogenic genes at different time points. Note that remains *REST* gene remains expressed throughout the entire differentiation process. **g**, PCA plot of samples used to study DNA methylation (MeDIP-seq). **h**, Heatmap represents top 50 differentially methylated peaks in astrocytes (day 50) versus iPSCs (day 0). Note that vast majority of genes are hypermethylated (49 out of 50) in astrocytes. **i**, Dot plot of DNA methylation state annotating peaks with highest variance for genes associated with NOTCH, JAK/STAT, mTOR, NCOR2 and others.

### Functional characterization of human astrocytes

Next, we performed a series of experiments to probe astrocyte-specific functions. It is well-known that astrocytes support neuronal structure and physiology. When co-culturing hPSC-astrocytes with glutamatergic cortical neurons for 8 days, multi-electrode array (MEA) experiments showed enhanced neuronal activity as compared to neuron-only cultures. This supportive effect of astrocytes provided to neurons was similarly detected when commercially available astrocytes were used as a control (iCell Astro from FUJIFILM CDI). By day 8, glutamatergic neurons co-cultured with astrocytes displayed robust electrical activities (**Fig. 4a**) as reflected by higher number of spikes (**Fig. 4b**; *p = 0*.*47*), higher average number of active electrodes (**Fig. 4c**, *p=0*.*0027*), and robustness of spike intervals across wells (**Fig. 4d**; *p=0*.*0117*) as compared to neuron-only cultures. In independent experiments, when astrocytes were co-cultured with motor neurons for 21 days, it was noted that astrocytes themselves showed remarkable structural plasticity and developed elaborate morphologies with GFAP^+^ processes (**Extended data Fig. 7b**). Such long projections of GFAP^+^ varicose structures were reported to be specific for human astrocytes^43^. In these co-culture experiments, motor neurons also developed more prominent puncta immunoreactive for the synaptic vesicle protein synapsin-1 (**Fig. 4e**). Furthermore, electrophysiological analysis of co-cultures showed higher network synchrony and reduced average inter-burst intervals (**Fig. 4f** and **4g;** *p = 0*.*18*) as a measure of synaptic maturation of the neuronal network. Next, to measure neuronal survival and neurite outgrowth in long-term cultures (21 days), hPSC-astrocytes were co-cultured with i3-neurons^44^, stably expressing green fluorescent protein (GFP) in the nucleus and red fluorescent protein (RFP) in the cytoplasm. By day 21, i3-neurons revealed a reduction in the number of GFP^+^ nuclei when cultured alone. In contrast, neuronal loss was prevented in co-cultures with iPSC-derived astrocytes (**Extended Data Fig. 7a**). To quantify neurite length, continuous live-cell imaging was performed using the IncuCyte system. Detectable from day 2 onward, RFP^+^ i3-neurons co-cultured with astrocytes developed neurites with increased length (**Fig. 4h** and **4i**). Next, since astrocytes are known to uptake extracellular neurotransmitters, we performed colorimetric analysis of L-glutamate levels after administration to the medium. Indeed, hPSC-astrocytes were capable of glutamate uptake comparable to controls (commercially obtained human and mouse astrocytes) (**Fig. 4j**). Next, since astrocytes play important neuro-immunological roles, we used an assay to confirm that hPSC-astrocytes could respond to microglia-secreted cytokines Il-1b, TNF-alpha and C1Q, by releasing complement component C3 (**Fig. 4k**). High-content calcium imaging revealed that day-30 hPSC-astrocytes can elicit transients when stimulated by KCL, ATP and L-glutamate (**Fig. 4l**). Astrocyte cultures were also found to exhibit slow spontaneous, calcium transients spreading across larger cell populations (**Supplementary Movie 1**) similar to control astrocytes (**Supplementary Movie 2)**. Lastly, glycogen accumulation as another important astrocyte characteristic was confirmed by Periodic Acid-Schiff staining showing a similar pattern in hPSC-astrocytes and controls (**Fig. 4m**).

**Fig. 4:**
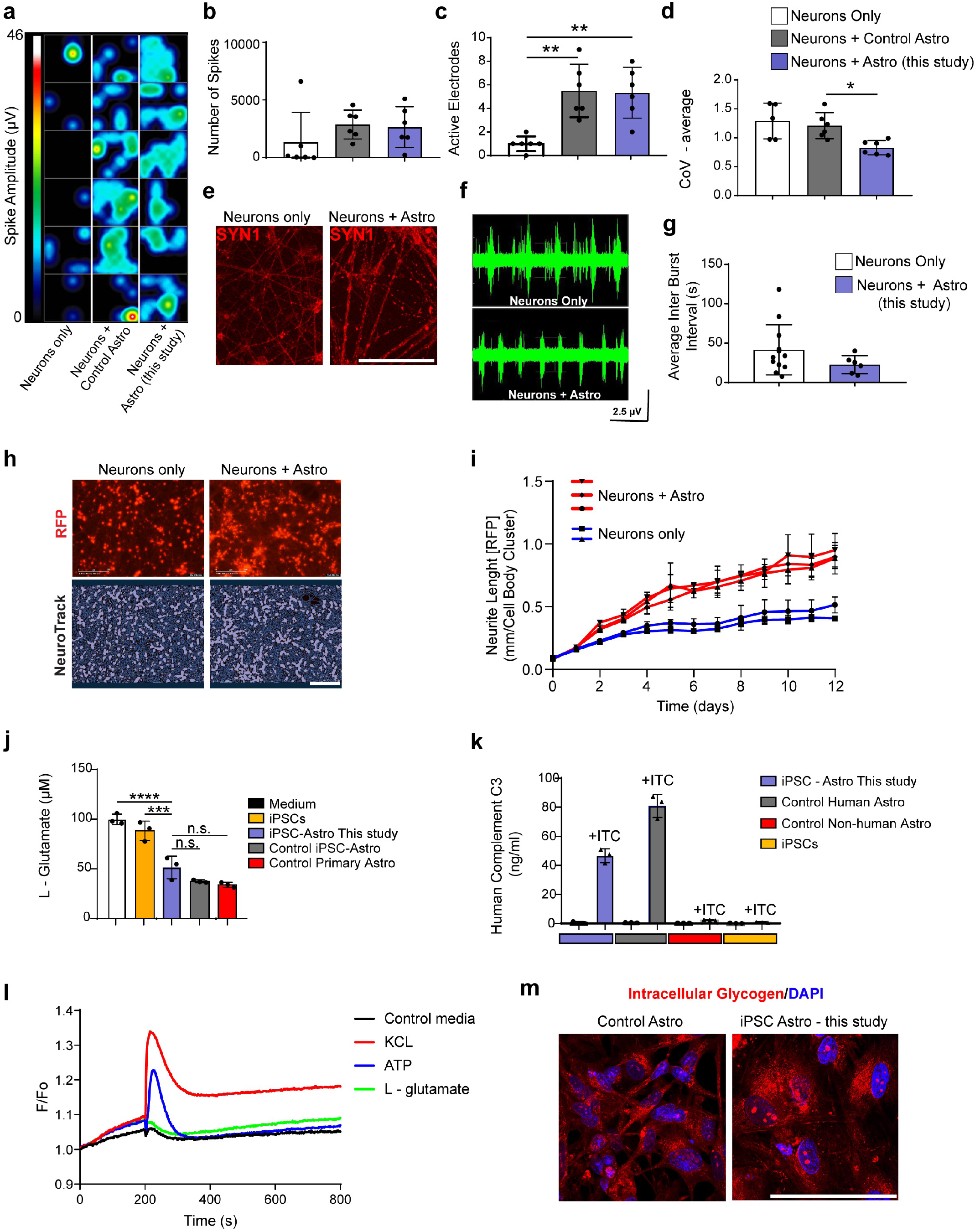
Functional characterization of hPSC derived astrocytes. **a**, Enhanced neuronal activity of glutamatergic cells upon co-culture with hPSC-astrocytes. Multi-electrode array (MEA) analysis was performed at day 8 of co-culture show. **b-d**. Quantification of spikes (b), active electrodes (c) and reduced coefficient of variation between recorded wells (d) in co-culture as compared to glutamatergic neurons only (n=6 wells per group). **e**, Immunoreactivity for synaptic marker synapsin-1 (SYN1) expressed by motor neurons co-cultured with astrocytes for 21 days versus neuron-only cultures. **f**, MEA recording (1 out of 48 electrodes represented) to visualize sporadic spikes in-between bursts in neuronal mono-culture versus higher synchrony in co-cultures. **g**, MEA analysis shows reduced inter burst interval when motor neurons are co-cultured with hPSC-astrocytes (unpaired t-test, p=0.18, nMN = 11; nMN + Astro = 6 wells). **h**, Video-microscopy and quantification of RFP^+^ neurite length over the course of 12 days in neuron-only versus neuron-astrocyte co-cultures. i3-neurons were co-cultured with astrocytes in a ratio of 1 astrocyte per 3 neurons (1:3). Images obtained with IncuCyte S3 live imaging system (each dot represents a well of a 24-well plate; and for each dot 4 images were taken and analyzed). Neurite length is normalized to cell body clusters. Note the clear separation of co-cultures from day 2 onward. **i**, Representative images of RFP^+^ i3-neurons and image mask of neurites and cell bodies (NeuroTrack software). **j**. Uptake of glutamate by iPSC-astrocytes is comparable to controls (primary mouse astrocytes and human iCell astrocytes (FUJIFILM CDI). **k**, Secretion of human complement C3 by hPSC-astrocytes into the culture medium after stimulation with inflammatory cytokines (Interleukin-1b, TNF-alpha and C1q) for 24 h. **l**, Calcium transients in iPSC-astrocyte cultures after stimulation with KCL, ATP and L-glutamate. **m**. Periodic acid-Schiff stain of intracellular glycogen in hPSC-astrocytes and controls (iCell Astro from FUJIFILM CDI). Scale bar, 100 µm (**e, h, m**).

### Translational utility of hPSC-astrocytes

Ideally, clinically relevant cell differentiation protocol should enable the large-scale biomanufacturing of human cells from hPSCs. To this end, we automated our scalable astrocyte differentiation protocol using the CompacT SelecT robotic platform (**Fig. 5a**). This approach enabled the production of nearly 500 million astrocytes in one single experimental run and astrocytes could be cryopreserved and used on-demand for various applications. For instance, to further characterize hPSC-astrocytes, we performed grafting experiments into mouse brains. Neonatal pups and adult mice received unilateral injections, either intraventricular and cortical, of iPSC-derived astrocytes and the brains were analyzed at 4 weeks and 8 weeks post-transplantation (**Fig. 5b-g)**. In all experimental groups, immunohistochemical analysis revealed strong immunoreactivity for the human cytoplasm marker (STEM121) at the injection sites as well different brains regions in the ipsi- and contralateral hemispheres. Migratory glial cells co-expressing human-specific GFAP (hGFAP; STEM123) and S100B were widely distributed and detectable in the cortex (**Fig. 5b**), corpus callosum (**Fig. 5c**), hippocampus (**Fig. 5d**), thalamus (**Fig. 5e**) and surrounding blood vessels within the brain parenchyma (**Fig. 5f**). Quantitative analysis of whole-brain sections showed that highest numbers of transplanted human astrocytes were detected in neonatal animals 8 weeks post-grafting (**Fig. 5g**). Together, these experiments demonstrate that hPSC-derived astrocytes engraft and survive *in vivo* for several weeks.

**Fig. 5:**
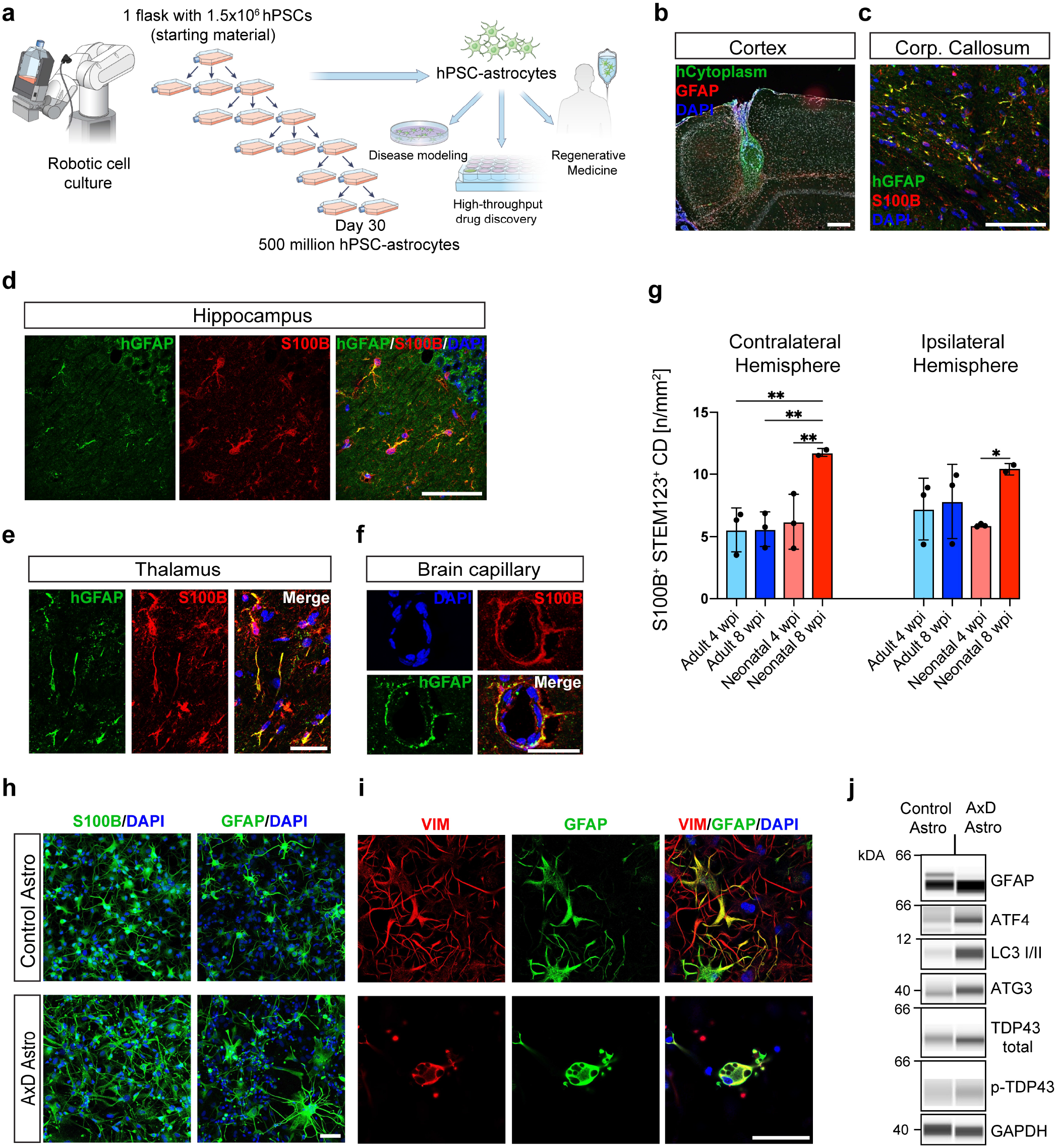
Robotic astrocyte differentiation enabling cell grafting and disease modeling. **a**, Schematic overview of robotic cell differentiation and large-scale astrocyte production using the CompacT SelecT (Sartorius). **b**, Human iPSC-astrocytes transplanted into the adult mouse cortex (4 weeks post-grafting) show strong immunoreactivity against human cytoplasmic marker STEM121 surrounded by glial scar formation (GFAP, red) **c**. Immunostaining for human-specific GFAP (STEM123) and S100B showing migratory cells in the corpus callosum. **d, e**, Immunostainings using human specific GFAP antibody (STEM123) depicts cells in thalamic and hippocampal regions indicating long-distance migration of astrocytes grafted into the cortex. **f**, Human GFAP expressing astrocytes surrounding a blood vessel in mouse brain parenchyma. **g**, Quantification of S100B and hGFAP co-labeled cells in the ipsi- and contralateral hemispheres of neonatal and adult mice at 4-and 8-weeks post-grafting (3 animals per group). **h**, iPSCs from Alexander disease (AxD) patient and an unaffected individual (control) were differentiated into S100B and GFAP expressing astrocytes. **i**, AxD astrocytes show typical GFAP^+^ protein aggregates and aberrant morphology. **j**, Western blot demonstrating strong upregulation of ER stress and autophagy markers. Total expression of TDP-43 expression was higher in AxD astrocytes versus control, whereas the phosphorylated form of TDP-43 was moderately higher in AxD.

Lastly, we established a cellular model for Alexander disease (AxD) by generating astrocytes from patient-derived iPSCs. Based on available information (Coriell), the AxD patient was heterozygous for a mutation in codon 239 of the GFAP gene with C-to-T transition at nucleotide 729 (729C>T) resulting in arg239-to-cys mutation *ARG239CYS* (R239C)^45^. Clinically, macrocephaly and regression were observed starting from 18-months of age and MRI scan showed white matter deterioration and ventricular cyst. The patient experienced first seizures at age of 3.5 years and died at 6 years of age and eventually the diagnosis was confirmed by autopsy. First, astrocytes expressing S100B and GFAP (day 50) were generated from control and AxD iPSCs (**Fig. 5h**). Confocal microscopy demonstrated aberrant morphology of AxD-astrocytes with prominent GFAP^+^ aggregates resembling Rosenthal fibers (**Fig. 5i** and **Extended Data Fig. 8**) consistent with previous reports^46,47^. Furthermore, AxD-astrocytes displayed abnormal unfolded protein stress response (UPR) as indicated by ATF4/ATG3 protein upregulation. Elevated expression levels of the autophagy marker LC3I/II, GFAP, and both total and phosphorylated TDP-43, which are associated with several neurodegenerative diseases, suggested the presence of additional pathophysiological mechanisms in AxD-astrocytes **(Fig. 5j**) that merit future research. Collectively, these findings showed that astrocytes derived with the iPSC differentiation method presented in this study are suitable for disease modeling.

## Conclusion

Derivation of human astrocytes from hPSCs has been a great challenge impeding their systematic utilization. Here, we developed a novel strategy that allows directed differentiation of hPSCs into RGCs and astrocytes without preceding neurogenesis. This rapid method is scalable and robust and generates large quantities of human astrocytes in the absence of serum and genetic manipulation. We envisage that streamlined access to iPSC-derived astrocytes will help to reduce the use of laboratory animals, which are currently the most popular source for astrocytes. Moreover, industrial-scale production of patient-and disease-specific astrocytes should help to deliver on the promise of the iPSC technology and leverage drug development, disease modeling, and personalized cell and gene therapies for various neuro-psychiatric illnesses.

## Methods

### Cell culture

hESC line (WA09), healthy donor iPSC lines (LiPSC-GR1.1 and NCRM5 from NIH Common Fund) and patient-derived Alexander disease iPSCs (GM16825, Coriell) were maintained under feeder-free conditions using Essential 8 (E8) Medium and vitronectin (VN)-coated plates (#A14700; Thermo Fisher Scientific). Cells were routinely passaged using 0.5 mM EDTA diluted in phosphate buffered saline (PBS) without calcium or magnesium (Thermo Fisher Scientific) when culture plates reached about 70–90% confluency, typically every 3 to 4 days. For the initial 24 h after cell passaging, E8 medium was supplemented with CEPT cocktail to optimize cell viability^20^.

### Astrocyte differentiation using Astro-1, Astro-2, and Astro-3 medium

Astro-1 medium was composed of DMEM/F12 medium supplemented with N2 (17502048, Gibco) and B27 without vitamin A (12587010, Gibco), 100 nM LDN193189 dihydrochloride (6053, Tocris), and human recombinant proteins PDGF-AA (221-AA, R&D Systems), JAGGED-1 (1277-JG, R&D Systems) DLL-1 (1818-DL, R&D Systems), ONCOSTATIN M (295-OM, R&D Systems), LIF (7734-LF, R&D Systems), CNTF (257-NT, R&D Systems) all at 10 ng/ml concentration. Astro-2 medium was composed of DMEM/F12 base medium supplemented with N2 (Gibco), B27 complete (17504044, Gibco), 1% chemically-defined lipid supplement (11905031, Gibco) and recombinant proteins JAGGED-1, DLL-1, ONCOSTATIN M, LIF, CNTF all at 10 ng/ml concentration (R&D Systems).

Astro-3 medium was composed of DMEM/F12 base medium supplemented with N2, B27 with vitamin A, 1% chemically-defined lipid concentrate (Gibco) and JAGGED-1, DLL-1, LIF, CNTF all at 10 ng/ml concentration, hNRG1/EGF domain 20 ng/ml (396-HB, R&D Systems), 2 µM forskolin (1099, Tocris), 200 nM phorbol-12 myristate-13 acetate (1201, Tocris), 40 ng/ml triiodothyronine T3 (6666, Tocris) and 200 µM ascorbic acid (4055, Tocris).

### Differentiation of hPSCs into RGCs and astrocytes

For RGC differentiation, hPSCs were detached using 0.5 mM EDTA and plated at 10.000 cell/cm^2^ density on VN-coated surface in E8 medium supplemented with CEPT (day -1). The following day medium was replaced with freshly prepared Astro-1. At day 3 cells were single cell dissociated using Accutase and 20.000 cell/cm^2^ was plated on VN-coated surface in Astro-1 medium supplemented with CEPT. Daily medium change was performed using fresh Astro-1 medium and cells appeared as neural rosettes by day 7.

For astrocyte differentiation, RGCs were maintained in Astro-1 medium until day 15, with daily medium changes and cell passaging performed on day 7, 11, and 14 using Accutase and at each passaging step 30,000 cells/cm^2^ were plated on VN-coated plates in Astro-1 medium supplemented with CEPT. On day 15, medium was replaced by Astro-2. Medium changes were performed daily with subculturing on day 18 and 23, and after cell counting 40.000 cells/cm^2^ were plated on VN-coated plates and maintained until day 30. At day 30, cells were either cryopreserved in Astro-2 medium supplemented with 10% DMSO and CEPT or further matured in Astro-3 medium until day 50.

For astrocyte maturation, day-30 astrocytes were plated on Geltrex (Thermo Fisher) at 50.000 cells/cm^2^ and maintained in Astro-3 medium until day 50. Astro-3 medium was composed of DMEM/F12 medium supplemented with N2, B27 with vitamin A, 1% chemically-defined lipid supplement (Gibco) and JAGGED-1, DLL-1, LIF, CNTF all at 10 ng/ml concentration, hNRG1/EGF domain 20 ng/ml, 2 µM forskolin, 200 nM phorbol-ester, 40 ng/ml triiodothyronine and 200 µM ascorbic acid. Cultures were passaged (1:2 ratio) on day 37 and 43, respectively. At day 50, cells were cryopreserved in Astro-3 medium supplemented with 10% DMSO and CEPT.

For robotic astrocyte differentiation, 1.75 million hPSCs were plated in T175 flasks and cultured in the CompacT SelecT (Sartorius) and Astro-1, Astro-2, and Astro-3 media were applied as described above. All media changes (Day 0-50) were performed by the robotic instrument throughout the cell differentiation process. Cell culture flasks were only briefly removed from the system for offline centrifugation to remove Accutase that was used for cell dissociation.

### Immunocytochemistry

Cells cultured in 6-, 24- or 96-well plates or 8 chamber slides (Ibidi) were fixed using 4% PFA diluted in PBS for 15 min and washed 3 times with PBS (5 min each). Blocking was performed with 4% donkey serum, 0.1% Triton-X in PBS for 1h at room temperature on a shaker. All primary antibodies used study were applied overnight at 4°C. Next day, cultures were washed 3 times with PBS and appropriate secondary antibodies were applied for 1h at room temperature. Fluorescence images were taken with the Leica DMi8 epifluorescence and Zeiss LSM 710 confocal microscopes using appropriate filters. Primary antibodies used for immunocytochemistry are as follows: mouse anti-PAX6 1:200 (561462, BD Biosciences), rabbit anti-FABP7 (BLBP) 1:200 (ABN14, EMD Millipore), rabbit anti-CD44 1:400 (ab157107, Abcam), rat anti-CD44 1:100 (A25528), mouse anti-VIMENTIN 1:500 (M0725, DAKO), rabbit anti-NFIA 1:250 (NBP-1-81406, Novus), rabbit anti-S100-beta 1:100 (ab52642, Abcam), mouse anti-TUJ1 1:1000 (801201, BioLegend), rabbit anti-GFAP 1:1000 (Z0334, Dako), rabbit anti-synapsin-1 1:500 (106-011, Synaptic Systems), rabbit anti-ASPM 1:100 (NB-100-227, Novus), rabbit anti-FAT1 1:100 (HPA023882 Millipore Sigma) and mouse anti-Pan-Neuronal Marker (PNM) 1:200 (MAB2300, EMD Millipore). The following secondary antibodies were used: donkey anti-mouse Alexa 568 (1:500, A-10037, Thermo Fisher) and donkey anti-rabbit Alexa Fluor 488 (1:500; A-21206, Thermo Fisher).

### Western blot

The Wes and Jes automated Western blotting systems (ProteinSimple) were used for quantitative analysis of protein expression as described previously^20,48^. All western blot data are displayed by lanes in virtual blot-like images. Briefly, cells were harvested by scraping, pelleted, washed with PBS, flash frozen using dry ice and stored at -20 °C until processed. Cell pellets were resuspended in RIPA buffer (Thermo Fisher) supplemented with halt protease inhibitor cocktail (Thermo Fisher Scientific) and lysed by sonication. Lysates were cleared of debris by centrifugation at 14,000g for 15 min and quantified using the BCA protein assay kit (Thermo Fisher Scientific). Lysates were diluted 1:4 with 1X sample buffer (ProteinSimple). Protein quantification was performed using the 12-230 kDa 25-lane plate (PS-MK15; ProteinSimple) in a Wes Capillary Western Blot analyzer according to the manufacturer’s recommendation. Protein quantification was done using the Compass software. Primary antibodies used are as follows: mouse anti-GAPDH 1:2.000 (sc25778, Santa Cruz), rabbit anti-PAX6 1:20 (901301, BioLegend), rabbit anti-FAT1 1:20 (HPA023882, Millipore Sigma), rabbit anti-ATF4 1:50 (D4B8, Cell Signaling), rabbit anti LC3 A/B 1:50 (D3U4C, Cell Signaling), rabbit anti-ATG3 1:50 (3415, Cell Signaling), rabbit anti-ATG5 1:50 (D5F5U, Cell Signaling), rabbit anti-TDP43 total 1:50 (89789, Cell Signaling) and rabbit anti-phospho-TDP43 1:50 (22309-1-AP, Thermo Fisher).

### Transmission Electron Microscopy (TEM)

Astrocytes differentiated from hPSC (day 30 and 50) and cultured in 6-well plates (1 million cells per well) and commercial hPSC-astrocytes (FUJIFILM CDI) were fixed by replacing the media with 2.5% glutaraldehyde in PBS (pH 7.2–7.4) for 60 minutes at room temperature. Further sample processing for electron microscopy and imaging (H7659) were performed by the Electron Microscopy Laboratory of Leidos Biomedical Research, NCI, Frederick, MD.

### Co-culturing astrocytes and neurons

For co-culture experiments, iPSC-derived glutamatergic or motor neurons (R1061, R1051; FUJIFILM CDI) were plated on Geltrex-coated plates (100,000 cells per cm^2^) and placed into the incubator (37 °C) for 45 mins to allow cells to attach. Then, 33,000 iPSC-astrocytes were added per cm^2^ to achieve a 1:3 ratio. The neuronal medium recommended by vendor (FUJIFILM CDI) was supplemented with LIF, CNTF (both 10 ng/ml) and 1% chemically-defined lipid supplement (Gibco). CEPT was included for the first 24 h. Co-cultures were maintained for up to 21 days prior to fixation with 4 % PFA and immunostaining.

### Multi-electrode array assay

Electrophysiology was performed using the Maestro platform (Axion Biosystems). hiPSC-derived glutamatergic neurons (R1062; FUJIFILM CDI) were plated at a density of ∼5 million neurons per cm^2^ in complete neuronal media recommended by the vendor as follows: BrainPhys Neuronal Medium (05790, STEMCELL Technologies) supplemented with iCell Neural Supplement B (M1029, FUJIFILM CDI) and iCell Nervous System Supplement (M1031, FUJIFILM CDI), N2 (Gibco), 10 μg/mL laminin (Thermo Fisher), 10 ng/ml LIF (R&D Systems), 10 ng/ml CNTF (R&D Systems) and 1% chemically defined lipid supplement (Gibco) and CEPT. 48-well plates with electrodes were coated with 0.1% polyethylenimine (PEI). hiPSC-derived motor neurons (FUJIFILM CDI) were plated at a density of ∼5 million neurons per cm^2^ in complete medium containing CEPT. For co-culture experiments with neurons, iPSC-astrocytes generated at NCATS were compared to iCell Astro (R1092, FUJIFILM CDI) that served as controls. Plating of astrocytes and neurons in 1:3 ratio is described above. 48-well MEA plates were coated with Geltrex (Gibco). Twenty-four hours post-plating, fresh medium was added and CEPT was removed and then 50% media changes were performed every 2-3 days. Electrophysiological recordings were performed daily for 10 mins.

### High-throughput calcium imaging

hPSC-derived astrocytes were plated onto Geltrex-coated 96-well plates with transparent bottom (Greiner) and maintained in Astro-3 media for 5-7 days with daily medium change prior to calcium imaging. Calcium imaging was performed on the FLIPR Penta high-content calcium imager (Molecular Devices) and on the day of the experiment, cultures were prepared according to the instructions detailed in the FLIPR Calcium Assay 6 kit (Molecular Devices). Briefly, cells were loaded using 100 μl of prepared loading buffer without removing media portion (100 μl) and incubated for 2 h at 37C. After incubation 96 well cell plate was transferred to FLIPR instrument and the experiment was initiated by taking 200 measurements of basal signal levels prior to supplementing cultures with either DMSO, L-glutamate (100 μM), ATP (30 μM) or KCl (65 mM), and 600 measurements upon transfer (read interval 1/s).

### Glutamate uptake assay

hPSC-derived astrocytes, commercial iCell astrocytes (FUJIFILM CDI), mouse astrocytes (ScienCell) and hPSCs (negative control) were plated onto Geltrex-coated 96-well plates (Corning) and maintained in Astro-3 medium (astrocytes), E8 medium (hPSCs) for 5-7 days with daily media changes prior to performing glutamate uptake assay. On the day of experiment, cells were first incubated for 30 min in Hank’s balanced salt solution (HBSS) buffer without calcium and magnesium (Gibco), prior to 3 h incubation with 100 μM L-glutamate in HBSS with calcium and magnesium (Gibco) to allow glutamate uptake by cells. After 3 h, medium/supernatants were analyzed with a colorimetric glutamate assay kit (Sigma-Aldrich) used according to the manufacturer’s instructions.

### Cytokine stimulation of astrocytes

hPSC-derived astrocytes, commercial hPSC-astrocytes (FUJIFILM CDI), mouse astrocytes (ScienCell) and hPSCs were plated onto Geltrex coated 96 well plates (Corning) and maintained in Astro-3 medium (astrocytes), E8 medium (hPSCs) for 5 – 7 days with daily medium changes prior to performing cytokine stimulation. On the day of the experiment cells were treated with 3 ng/ml IL-1α (SRP3310, Millipore Sigma), 30 ng/ml tumor necrosis factor (8902SF, Cell Signaling Tech) and 400 ng/ml C1q (MBS143105, MyBioSource) for 24 h. The following day medium was isolated, spun down to remove debris, and human complement C3 levels were measured using the Human Complement C3 ELISA Kit (ab108823, Abcam) following the manufacturer’s instructions.

### Glycogen staining (Periodic Acid-Schiff)

hPSC-derived astrocytes and commercial astrocytes (FUJIFILM CDI) were washed once with ice-cold PBS and fixed with ice-cold methanol for 5 min. After fixation, the Periodic Acid-Schiff staining kit (Millipore Sigma, 101646) procedure was followed as described by manufacturer. Autofluorescence emitted by PAS-labeled granules was captured by using the Leica DMi8 epifluorescence microscope at 568 nm wavelength.

### Cell transplantation

Adult male and female C57Bl/6J mice were purchased from The Jackson Laboratory (Maine, USA) at 8 weeks of age and grouped housed in same gender in OptiMice ventilated racks. Mice were acclimated to the vivarium for a week prior to injections. For neonatal injections, time pregnant C57Bl6/J mice (E13 – E15) were purchased from The Jackson Laboratory. Upon receipt mice were single housed in OptiMice ventilated racks. Pups were weaned at 21 days and housed by gender and treatment group and kept in 12/12 light/dark cycle. The room temperature was maintained between 20 and 23°C with a relative humidity around 50%. Chow and water were provided ad libitum.

On the day of cell grafting, media was aspirated from the T25 flasks with iPSC-derived astrocytes (day 30 of differentiation) and 2 ml of Accutase (room temperature) was added per flask and incubated for 7 min, after which the cells were monitored by microscopy every 2-3 minutes until rounding of the cells was observed. Astrocytes were then pooled in sterile conical tubes and centrifuged at 300g at room temperature for 3 min. Cell pellets were gently resuspended in 1-5 ml DPBS. Cell numbers were counted with Countess Counter (Thermo Fisher) with Trypan Blue staining. Live cell numbers were calculated, and the cell suspension was supplemented with CEPT. After resuspension, cells were kept on ice and made available for injection. 3μl of suspension/50.000 cells per hemisphere unilaterally was used.

To perform intracerebroventricular (ICV) injections into newborn mice (postnatal day 1), animals were exposed to cryoanesthesia and injected with a dose volume of 1μl/g solution. A micro-liter calibrated sterilized glass micropipette was used that was attached to a 10 μl Hamilton syringe. The needle tip was adjusted to correct length for a 2 mm penetration into the skull. The immobilized mouse was firmly grasped by the skin behind the head. A fiber optic light was used to illuminate relevant anatomical structures used as a guide. The needle was penetrated, perpendicularly, 2 mm into the skull, for ICV at a location approximately 0.25 mm lateral to the sagittal suture and 0.50-0.75 mm rostral to the neonatal coronary suture. Dosing solution was dispensed slowly (about 1μl/sec). Once the dose was administered, the needle was slowly removed (about 0.5 mm/sec) to prevent back flow.

For grafting into adult mice, animals were placed in an induction chamber to receive anesthesia using 2% isoflurane in the air mixture. The fur of the scalp and anterior back was clipped, and the animal was placed back into the induction chamber. The animal was then placed in a stereotactic apparatus such that the 180 ear bars were in the ear canals and the incisors were in the tooth bar of the mouse adapter. Surgical planes of anesthesia were maintained with 2% isoflurane by a nose cone fitted to the stereotaxic instrument. The scalp was prepared for surgery by cleaning the clipped injection site with Povidone iodine and rinsed with 70% ethanol. Mice were checked for a withdrawal reflex by pinching the hind foot to assess for anesthetic depth before an incision was made. A 1-1.5 cm slightly off-center incision was made in the scalp. The subcutaneous tissue and periosteum were scraped from the skull with sterile cotton tipped applicators. The guide cannula attached to stereotaxic arm was positioned over Bregma and this coordinate is considered the zero point. The guide cannula was moved to the appropriate anterior/posterior (−1.9) and medial/lateral (+/- 0.3) coordinates. Mice were infused with cells bilaterally in the cortex at a depth of (−2.5) with 3ul on each side, at a rate of 0.3 μl/min with a 4 min rest period following injection. Overall, one adult mouse and 3 pups died post-injection.

For brain collection, animals were randomly split into two groups for collections at 2 time points after engraftment: 1) 4 weeks post grafting; 2) 8 weeks post grafting. Mice were deeply anesthetized with pentobarbital and monitored for loss of reflexes until all the responses to external stimuli cease (verified by a toe pinch). The abdominal cavity was opened, followed by opening of the chest cavity. A blunt-end perfusion needle was inserted into the left ventricle and forwarded toward the ascending aorta. Immediately after, a small incision was made on the right atria. The perfusion needle was connected to an automated perfusion pump through a catheter system for a continuous administration of ambient temperature phosphate-buffered saline (PBS; 20 mM, pH 7.4 at room temperature). Approximately 40 ml of PBS was perfused at a rate of 10 ml/min. Saline flush was followed by perfusion with freshly prepared phosphate buffered 4% PFA. Brains were harvested and post-fixed overnight in freshly prepared phosphate buffered 4% PFA then transferred to 15% sucrose/PBS solution.

For immunohistochemistry, cryosectioned brains were stained to detect expression of human cytoplasmic marker STEM121 (TaKaRa, cat. Y40410) and human-specific GFAP marker STEM123 (TaKaRa, cat. Y40420). As general astrocyte markers GFAP (DAKO; z0334) and S100B (Abcam; ab52642) were used.

For quantitative image analysis, four different ROIs were manually drawn in Image J and either concentrated on the cerebral cortex around the graft injection site or delineated the entire hemisphere. A bilevel black and white DAPI mask was stored for every ROI and image. The inverted DAPI mask was mathematically subtracted from the S100B original image channel. A mask of S100B positive nuclei was stored for each image and ROI. The mask of S100B positive nuclei was mathematically added to the original S100B image. This procedure allowed for extraction of the entire shape of cells around the DAPI^+^ and S100B^+^ nucleus as far as labeled. This approach guaranteed that only structures that can be measured are those directly attached, hence, segmented processes were not included. A mask reflecting this core shape of S100B^+^ astrocytes was saved for each image and ROI. The inverted S100B cell mask was subtracted from the original STEM123 image and signal above threshold and size restrictions overlapping with S100B^+^ morphology was counted using restrictions and “fill holes” and “pixel connect” option to assure that for every IR area one object is counted (not to bias objects density). All cell transplantation experiments were performed by PsychoGenics (Paramus, NJ, USA) as part of a service agreement.

### Bulk RNA-seq analysis

Bulk RNA-Seq samples (iPSCs-day 0, IPSC derived RGCs-day 7 (Astro-1), NSCs day 7 (dSMADi), day 14, day 21, and iPSC-astrocytes from day 30 and day 50) were preprocessed with a standard pipeline that can be viewed at https://github.com/cemalley/Jovanovic_methods. Software used in preprocessing included fastqc 0.11.9, STAR 2.7.8a, trimmomatic 0.39, htseq 3.7.3 (http://www.bioinformatics.babraham.ac.uk/projects/fastqc no publication for fastqc only this website, https://academic.oup.com/bioinformatics/article/29/1/15/272537, https://academic.oup.com/bioinformatics/article/30/15/2114/2390096, https://academic.oup.com/bioinformatics/article/31/2/166/2366196). Analysis was performed in R 4.0.3 (R Core Team 2021 https://www.R-project.org/). Samples were combined with RUVSeq batch correction method RUVg using a set of housekeeping genes (https://www.nature.com/articles/nbt.2931, gene list: https://bmcmedgenomics.biomedcentral.com/articles/10.1186/s12920-019-0538-z#Tab3).The UCSC Cell Browser cortex development dataset was mined to find unique differentially expressed genes among the radial glia cell subpopulations RGdiv1, oRG, or Pan-RG (https://science.sciencemag.org/content/358/6368/1318.long, https://cells.ucsc.edu/?ds=cortex-dev). DESeq2 was used to merge and normalize samples with the median-of-ratios method (https://genomebiology.biomedcentral.com/articles/10.1186/s13059-014-0550-8). Normalized counts from the resulting R object were row Z-score transformed and scaled before plotting a subset of the DE RG genes, key pluripotency and astrocyte markers in a heatmap using ComplexHeatmap 2.6.2 (https://academic.oup.com/bioinformatics/article/32/18/2847/1743594). Several public RNA-Seq datasets were obtained from the Sequence Read Archive and merged with the day 30 and day 50 iPSC-astrocyte samples (Ext. data figure 4): PRJNA412090 or GSE104232 (Tchieu et al., https://www.ncbi.nlm.nih.gov/bioproject/PRJNA412090); PRJNA382448 or GSE97619 (Santos et al., https://www.ncbi.nlm.nih.gov/bioproject/PRJNA382448); PRJNA383243 or GSE97904 (Tcw et al. https://www.ncbi.nlm.nih.gov/bioproject/PRJNA383243); and PRJNA297760 or GSE73721 (Zhang et al. https://www.ncbi.nlm.nih.gov/bioproject/PRJNA297760). Merging was done with the same RUVSeq RUVg method as previous. Principal component analysis plots were created in R with base function prcomp. Gene set enrichment was performed using the top 100 DE genes per contrast and input to Enrichr, querying the Gene Ontology 2019 or ARCHS4 Tissues databases (Fig.2c).

### ATAC-seq and MeDIP-seq analysis

#### Raw Data

ATAC-seq and MeDIP-seq experiment were performed by Active Motif (Carlsbad, CA, USA) and initial analyses provided included the following: 1. Sequencing and mapping paired-end Illumina sequencing reads to the reference human hg38 genome using BWA; 2. Alignment and peak calling statistics (using MACS2); 3. Generating summary statistics across samples comparisons; 4. Annotation of all intervals (peak regions); 5. Gene-centric rollup of the intervals table; 6. A top table comparing peak region metrics between different samples; 7. A UCSC custom tracks BED file; and 8. A signal map bigwig URL file for viewing in the UCSC viewer.

#### Modified differential peak analysis

Active Motif included all identified peaks in MACS2. To conservatively assess differential accessibility or methylation, only peaks supported by at least two biological replicates were retained for downstream analysis. DESeq-2 was then run on the filtered peaks (separately for ATAC-seq and MeDIP-seq) using default parameters.

For MeDIP-seq analysis reference genome for human hg38 was generated in BWA, paired-end fastq files aligned to the reference and counts generated using Rsubread with featureCounts (specifying paired-ends). DESeq2 was then used to normalize counts using the “median by ratios” method and used to run differential peak analysis for all pairwise comparisons (iPSC day 0 vs RGCs day 7; iPSC Day 0 vs astrocytes D30 and astrocytes D50; RGC Day 7 vs astrocytes D30 and astrocytes D50).

#### Overrepresentation analysis

After performing differential peak analysis, peaks were retained if their adjusted p-value was less than 0.05, the peaks ranked by adjusted p-value and overrepresentation analysis was performed using all Gene Ontology (GO) gene sets and Kyoto Encyclopedia of Genes and Genomes (KEGG) 2019 with the R library, clusterProfiler using Benjamini Hochberg correction and an adjusted p-value cut-off of 0.01.

#### Principle Component Analysis (PCA)

For both ATAC-seq and MeDIP-seq data, PCA was performed to determine each peak’s variability across samples and genome-wide. The purpose behind performing this analysis was to determine which peak drives the variance across a given gene locus. For some gene loci, there were several identified ATAC-seq and MeDIP-seq peaks; however, it was not clear which peak might be driving differential accessibility or methylation. To perform this analysis, normalized count data was used in the prcomp function in R to perform PCA. For each gene, the peak with the largest variance was extracted from the PCA. Only peaks within 1,000 base pairs of an annotated gene were included in this analysis (which excludes enhancer elements that are far up-or downstream of a given gene).

The pre-filtered peak per gene with the highest variance was used to build dot plots for a targeted list of genes. All samples and normalized counts were used to calculate the interquartile range (IQR) for each gene (peak), separately. For each gene, peak and sample, the normalized count data was specified by the following criterion: 1. The median of the normalized counts was less than 0.25 quartile: Inaccessible (hypo-methlyated); 2. The median of the normalized counts was greater than 0.75 quartile: Hyper-accessible (hyper-methylated); or 3. The median of the normalized counts was between 0.25 and 0.75 quartiles: Mid-accessible (mid-methylated). The above criterion was used to generate the dot plots shown in Fig. 3b, i.

### Data presentation and statistical analysis

Data are presented as the mean ± s.d. unless otherwise stated in the respective figure legends. Statistical analyses (GraphPad Prism) were performed using different tests as appropriate and as described in figure legends.

## Supporting information

Ext Data Fig 1

Ext Data Fig 2

Ext Data Fig 3

Ext Data Fig 4

Ext Data Fig 5

Ext Data Fig 6

Ext Data Fig 7

Ext Data Fig 8

## Extended Data Figure Legends

**Extended Data Fig. 1: Consistent differentiation in additional hESC and iPSC line**

**a**,**b**, Widespread expression of PAX6, FABP7, and SOX9 in neural rosettes by day 7 in hESCs (WA09) and iPSCs (LiPSC-GR 1.1) differentiated with Astro-1. **c**, Representative images showing nuclear NFIA expression in most cells. **d**, Astrocytes express GFAP and VIM by day 50. Scale bars, 100 µm.

**Extended Data Fig. 2: Characterization of RGCs and astrocytes**

**a**, *BIRC5* expression in RGCs as detected by *in situ* hybridization (RNA-scope) after Astro-1 treatment or dSMADi (day 7). **b**, Quantification of *BIRC5* mRNA transcripts normalized to DAPI^+^ nuclei (n = 4 fields of view per group; *unpaired t-test *p=0*.*0115*). **c**, Representative images of ASPM- and PAX6-labeled cells at day 7 of differentiation. **d**. Quantification of mitotic spindle protein ASPM normalized to PAX6^+^ cells after differentiation with Astro-1 or dSMADi (n = 6 fields of view, *Unpaired t-test p=0*.*24)*. **e**, Immunostaining showing FAT1 expression by neural rosettes. **f**, Expression of S100B in differentiating cells (day 15). **g**, Quantification of S100B positive cells in two different hPSC lines normalized to total cells at day 15 (n = 4). **h**, Diffuse immunoreactivity for GFAP in immature astrocytes (day 30). **i**. Stellate morphology of GFAP^+^ astrocytes derived from LiPSC-GR1.1 (day 50). Scale bars, 100 µm.

**Extended Data Fig. 3: Ultrastructural analysis of astrocytes**

**a**, Overview images showing typical stellate morphology of iPSC-derived astrocytes. **b**, Astrocytes develop prominent tight junctions (red arrowheads). **c**, Representative examples of astrocytic processes (red dashed lines) with abundant intermediate filaments. Scale bars, 10 µm (**a**), 0.5 µm (**b, c**).

**Extended Data Fig. 4: Comparative transcriptomic analysis of hPSC-astrocytes**

**a**. PCA plot of iPSCs and astrocytes generated in the present study (serum-free and 2% FBS) and comparison to previous studies after performing batch correction. **b**, Hierarchical clustering based on the top 100 genes expressed in primary human fetal astrocytes according to ref. 31. Note that genes regulating histone proteins are prominently upregulated in iPSC-derived astrocytes (day 30) after addition of 2% FBS to Astro-2 medium. Note that these histone proteins are expressed in fetal brain-derived astrocytes (red boxed area). **c**, Gene enrichment analysis of the prominent histone gene clusters that was induced by 2% FBS (red boxed area in **b**).

**Extended Data Fig. 5: Targeted analysis of epigenetic changes within astrocyte gene markers in time-course of differentiation**

**a**. Chromatin accessibility in hPSCs (day 0) and astrocytes (day 30 and 50) considering the top 300 genes expressed by fetal and adult astrocytes (ref. 31). Heatmaps show gene peaks with highest variance for chromatin accessibility. Dynamic chromatin changes identify two dominant gene clusters (upper panel showing 31 genes and lower panel shows 79 genes) with open chromatin. **b**-**e**, Enrichr analysis of identified genes (*GO Biological process* and *ARCHS4 Tissues*) indicate biological processes such as cell motility, cell migration, and cell cycle.

**f**, Manhattan plot displaying differential methylation peaks with strongest difference between immature GFAP^-^ (day 30) and mature astrocytes GFAP^+^ (day50). *NCOR2* hypomethylation in mature astrocytes representing the most significant hit.

**Extended Data Fig. 6: Mapping of time-dependent chromatin accessibility of neurogenic and gliogenic genes**

**a, b**, Interactive Genome Viewer (IGV) plots showing chromatin accessibility of transcription factors that promote neurogenesis (*NEUROG2 and NEUROG3*). Note that chromatin regions are inaccessible for both transcription factors at day 30 and 50. **c, d**, IGV plots of chromatin accessibility for glial transcription factor NFIA and NFIB. Note that chromatin regions for both transcription factors are accessible at day 30 and 50.

**Extended Data Fig. 7: Functional characterization of astrocytes in co-culture with iPSC-derived neurons**

**a**, Microscopic images of i3-neurons expressing cytosolic RFP and nuclear GFP at day 21. See the difference between neuron-only versus co-culturing neurons with astrocytes. **b**, Overview image and photomontage of confocal images showing GFAP^+^ iPSC-astrocytes co-cultured with iPSC-derived motor neurons for 21 days (PNM = pan-neuronal marker). Note the long projection and ramifications of GFAP^+^ processes (white arrowheads). Scale bar, 100 µm.

**Extended Data Fig. 8: Confocal microscopic analysis of iPSC astrocytes from AxD**

Confocal z-stack images (1 µm sections) show GFAP^+^ aggregates and aberrant astrocyte morphology in AxD astrocytes (**a, b**) but not in iPSC-astrocytes from an unaffected individual (**c, d**). Scale bar, 50 µm.

**Extended Data Movie 1: Slow spontaneous calcium transients in iPSC-astrocytes**

Video-microscopy of iPSC-derived astrocytes (day 50) loaded with calcium dye (FLIPR Calcium 6 kit). Note the spreading of calcium transients across the cell culture. Images taken every 0.6 seconds.

**Extended Data Movie 2: Slow spontaneous calcium transients in control astrocytes**

Video-microscopy of iCell astrocytes (R1092; FUJIFILM CDI) loaded with calcium dye (FLIPR Calcium 6 kit). Note that these control astrocytes behave similarly to the iPSC-astrocytes shown in Extended Data Movie 1. Images taken every 0.6 seconds.

## Acknowledgments

We thank H. Hong, C. Sen, P. Francis, T. Deng, J. Slamecka, J. Freilino, T. Voss, M. Iannotti, C. Pepper Bonney, Y. Gedik, D. Ngan, A. Rossoshek, and A. Knebel for their support throughout this work. We are grateful to A. Hoofring from the NIH Medical Arts Design Section for art designs, K. Nagashima from the Electron Microscopy Laboratory at National Cancer Institute (NCI) for electron microscopy images, M. Hirst and D. Galbraith from Rancho BioSciences for bioinformatics support and Daniel Havas from PsychoGenics for confocal microscopy and quantification of grafted cells. We also gratefully acknowledge funding from the Regenerative Medicine Program (RMP) of the NIH Common Fund and in part by the intramural research program of the National Center for Advancing Translational Sciences (NCATS), NIH. The funders had no role in study design, data collection, and analysis; decision to publish; or preparation of the manuscript.

## Contributions

V.M.J. and I.S. conceived the project. Experiments: V.M.J., C.A.T., S.R., P.-H.C., E.B, P.O., J.C.M and S.M. Data analysis and discussions: V.M.J, C.M., C.A.T., M.W., A.S., I.S. Manuscript writing: V.M.J and I.S.

