## Supplementary figures and images for "Directed Differentiation of Human Pluripotent Stem Cells into Radial Glia and Astrocytes Bypasses Neurogenesis"

### Ext Data Fig 1

#### Extended Data Fig. 1 (Jovanovic et al.)

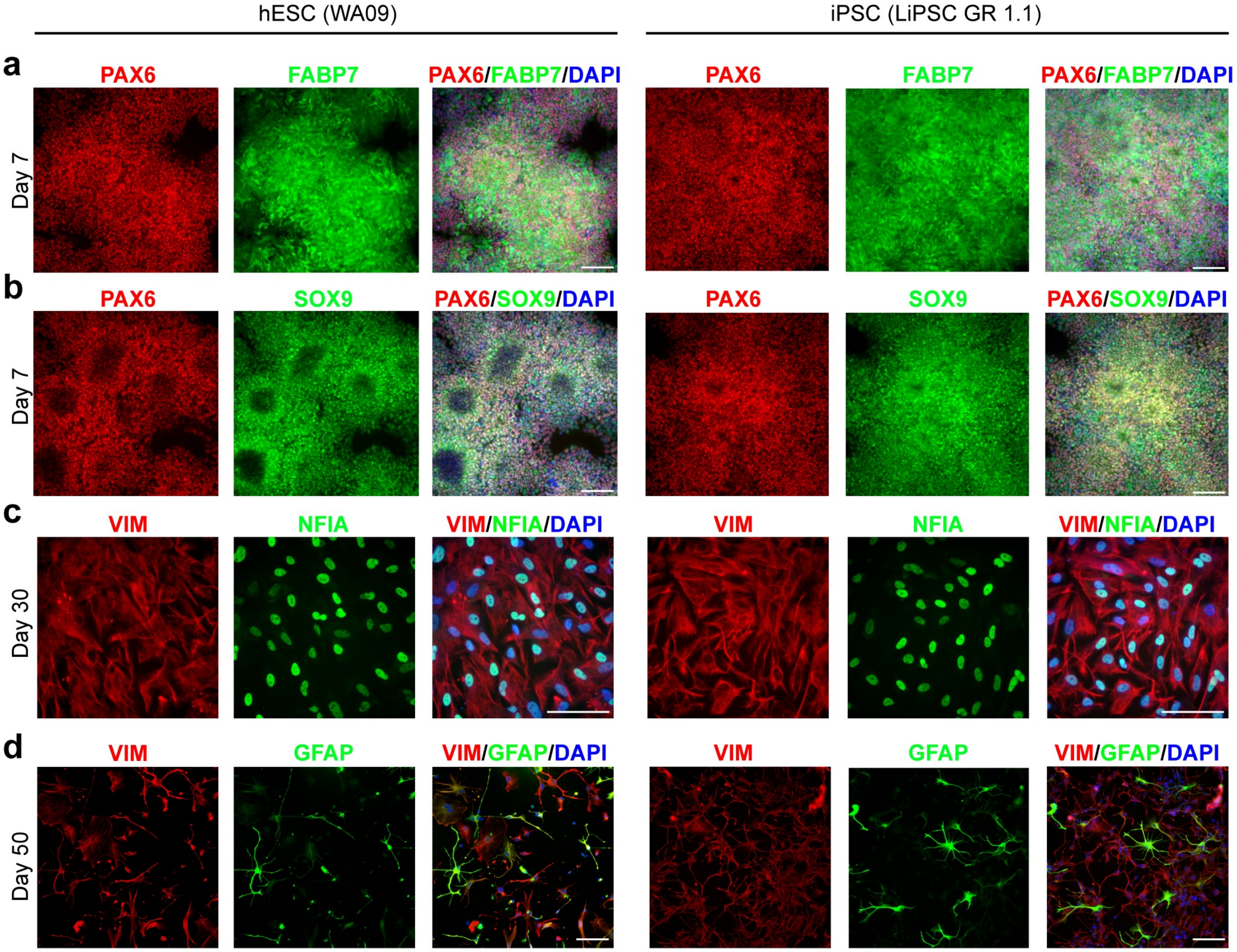

### Ext Data Fig 2

**Extended Data Fig. 2 (Jovanovic et al.)**

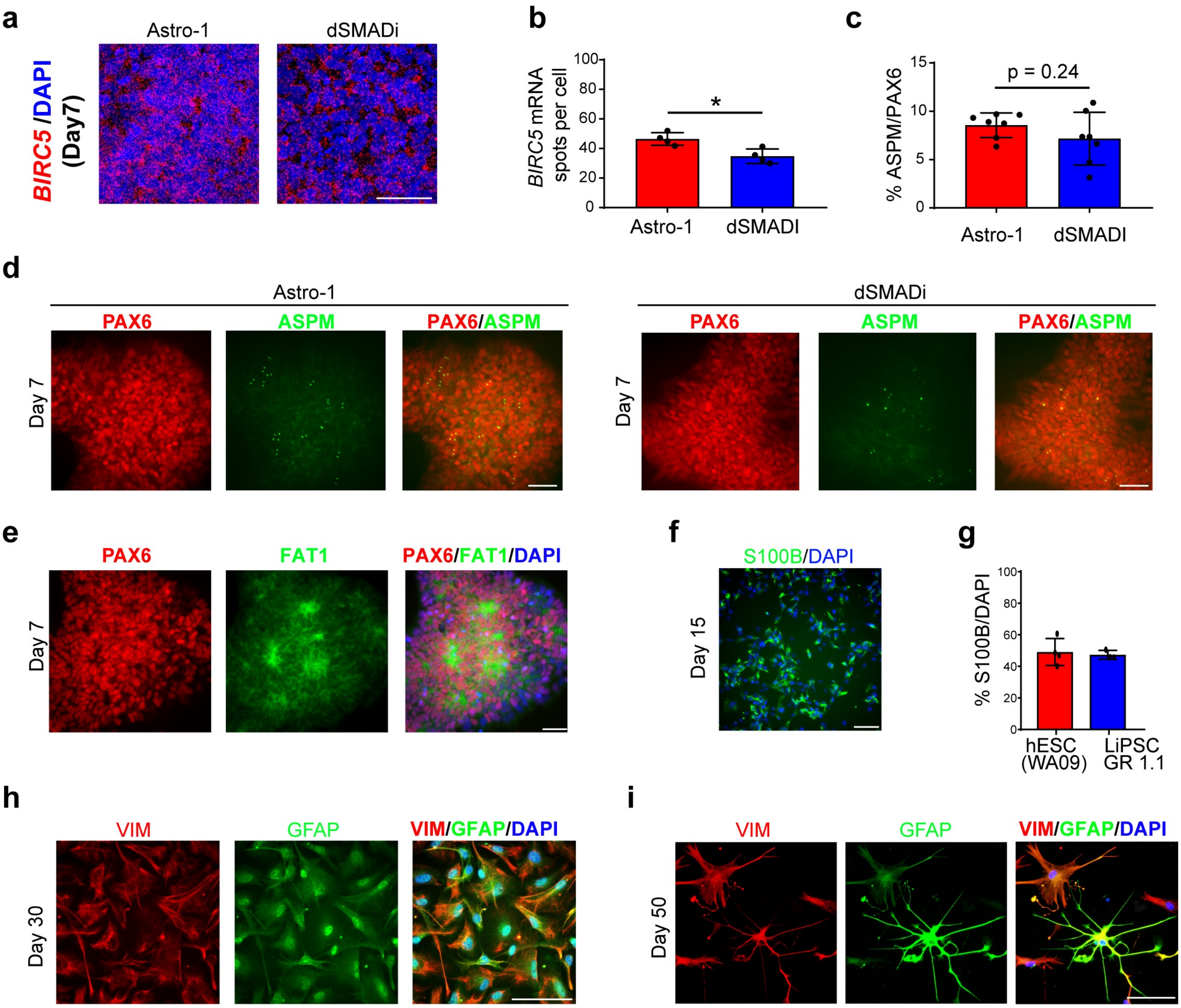

### Ext Data Fig 3

**Extended Data Fig. 3 (Jovanovic et al.)**

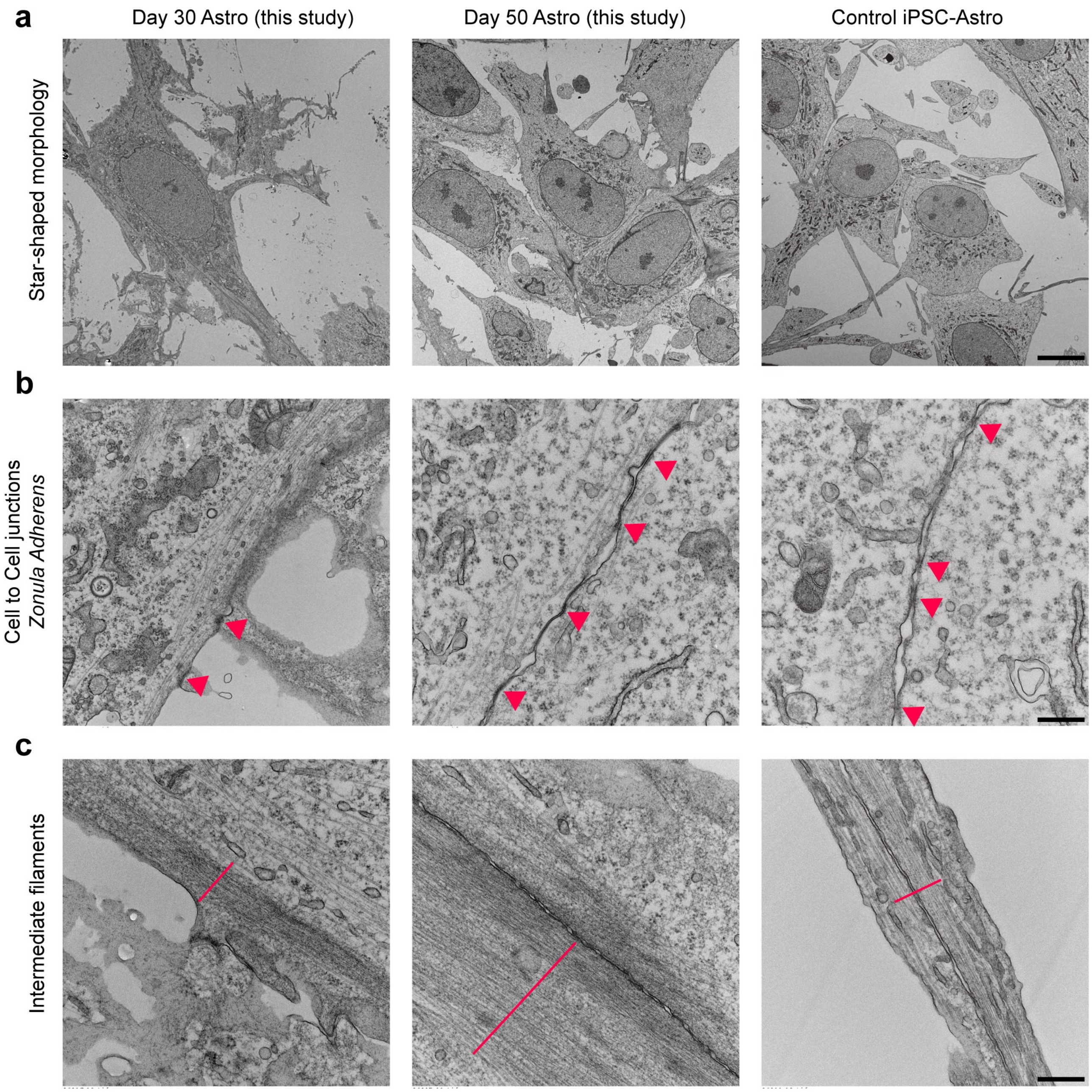

### Ext Data Fig 4

#### Extended Data Fig. 4 (Jovanovic et al.)

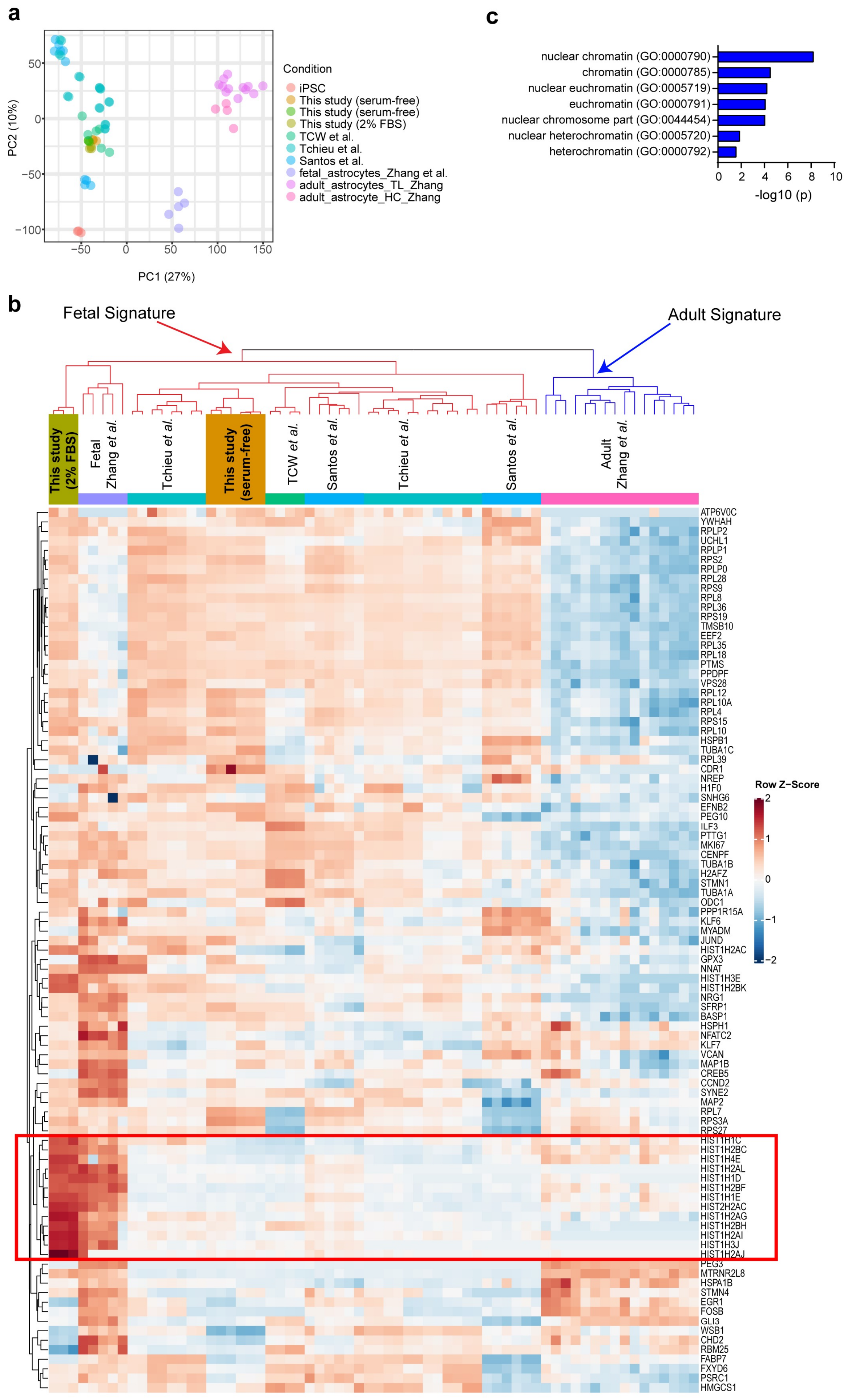

### Ext Data Fig 5

#### Extended Data Fig. 5 (Jovanovic et al.)

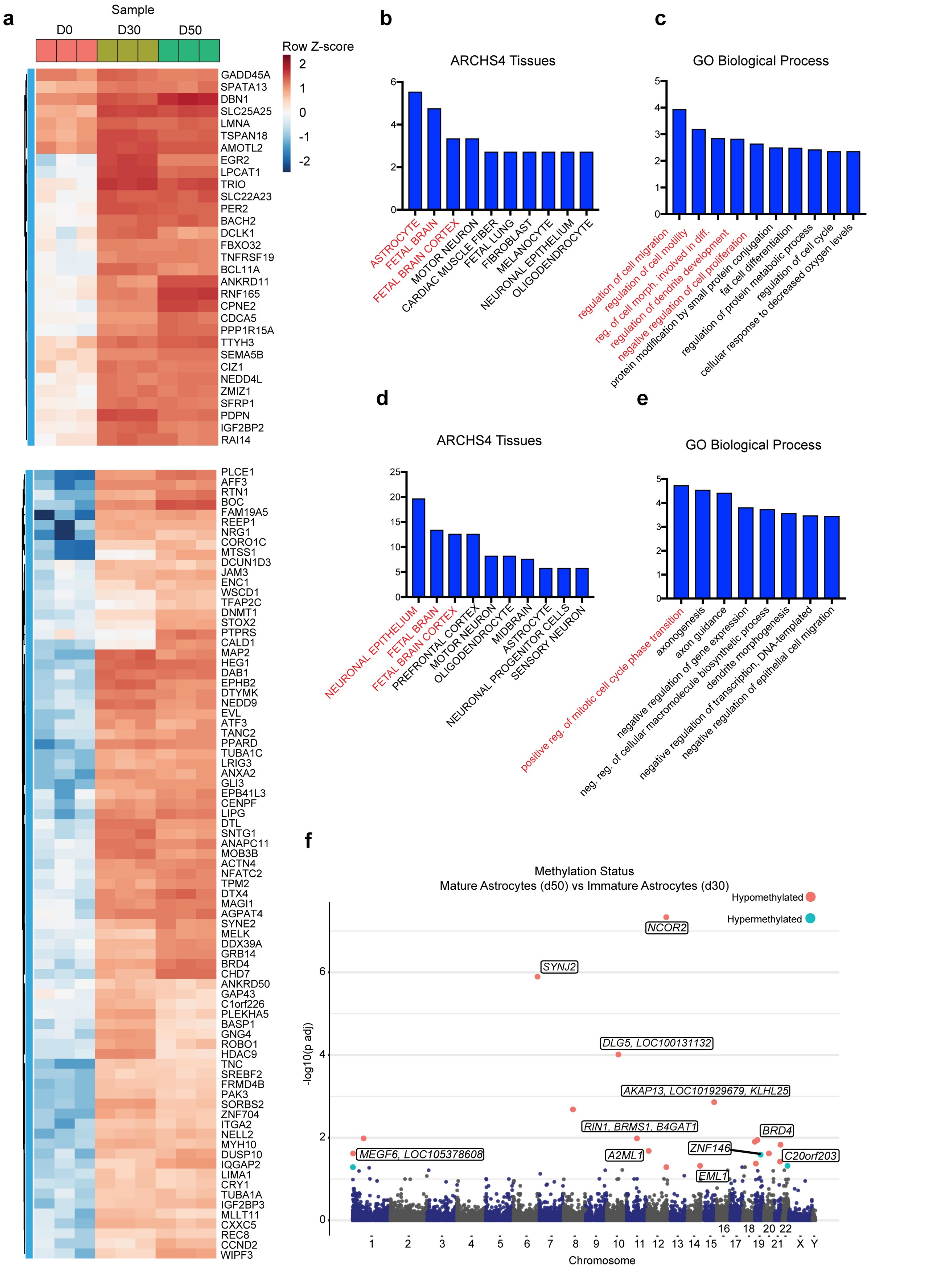

### Ext Data Fig 6

**Extended Data Fig. 6 (Jovanovic et al.)**

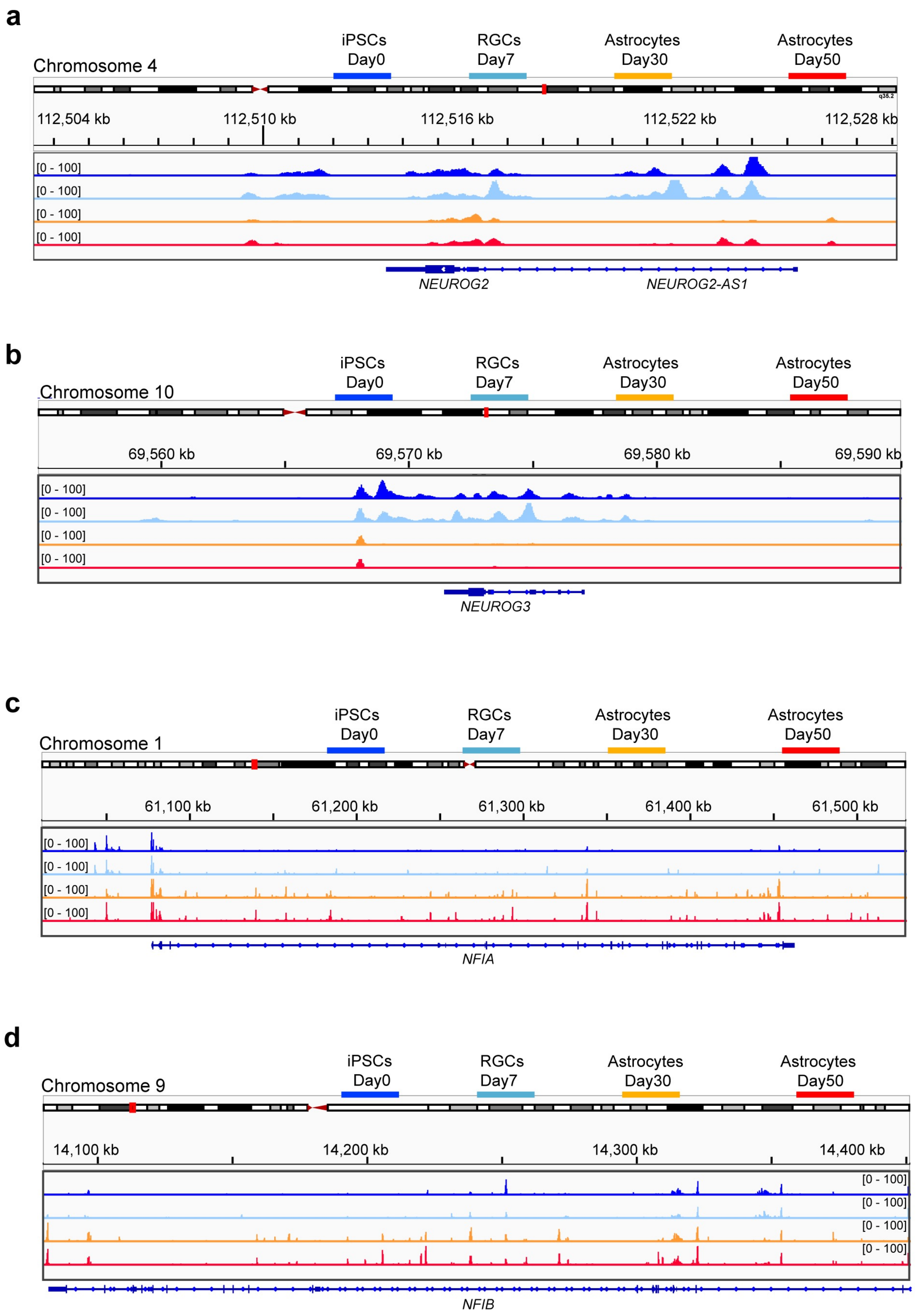

### Ext Data Fig 7

**Extended Data Fig. 7 (Jovanovic et al.)**

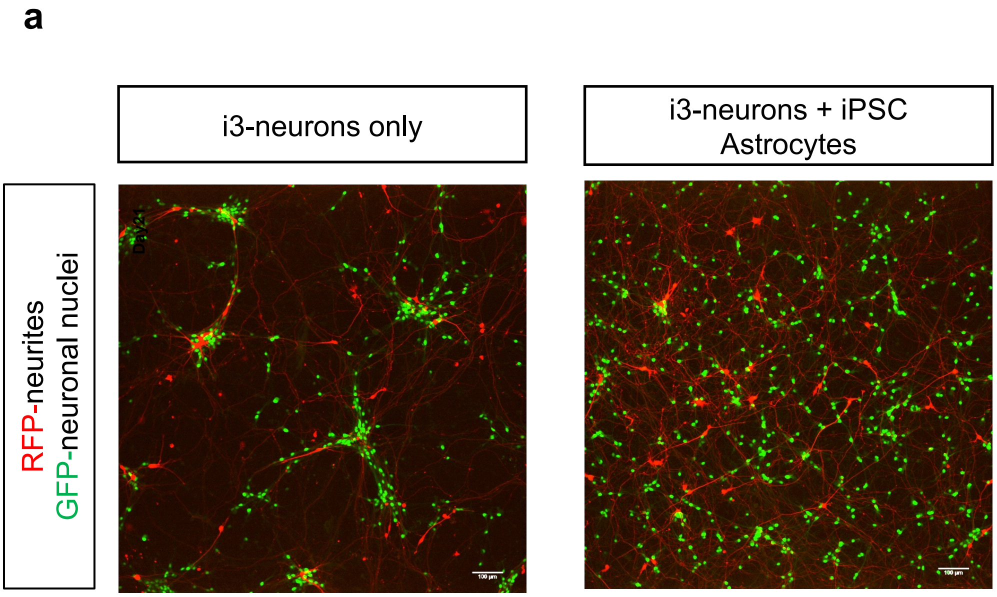


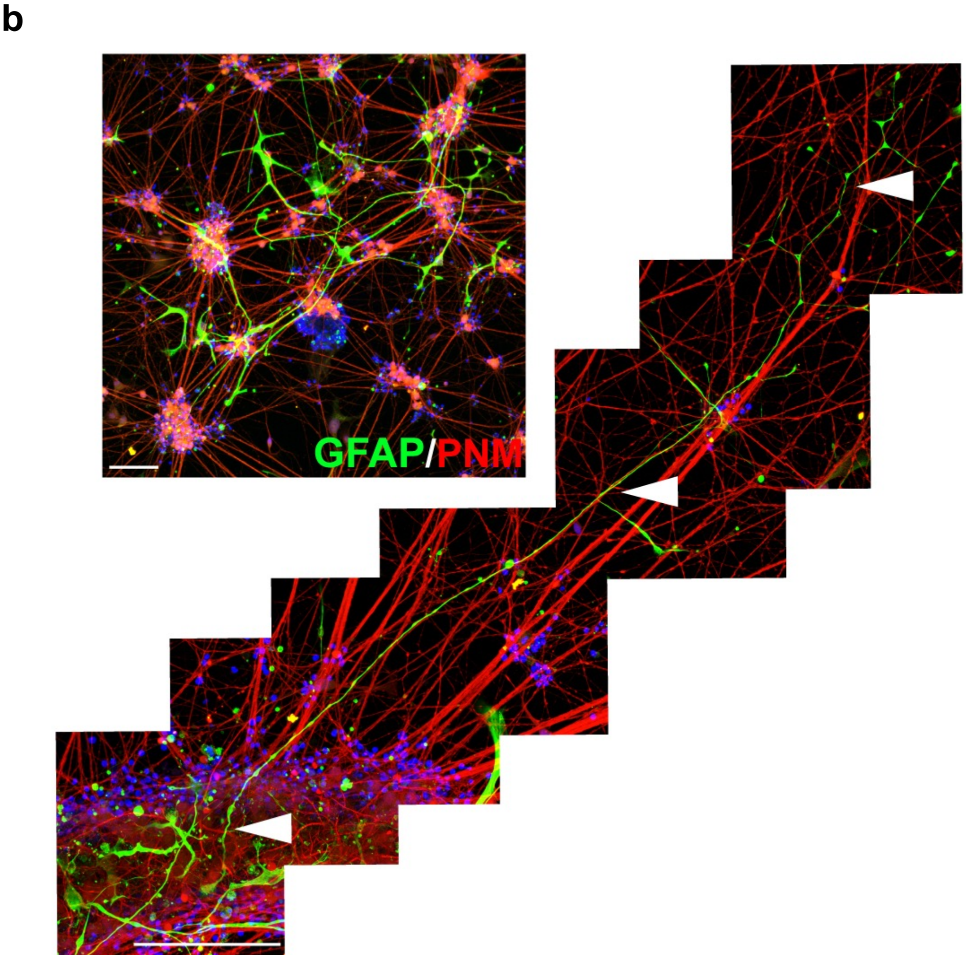

### Ext Data Fig 8

**Extended Data Fig. 8 (Jovanovic et al.)**

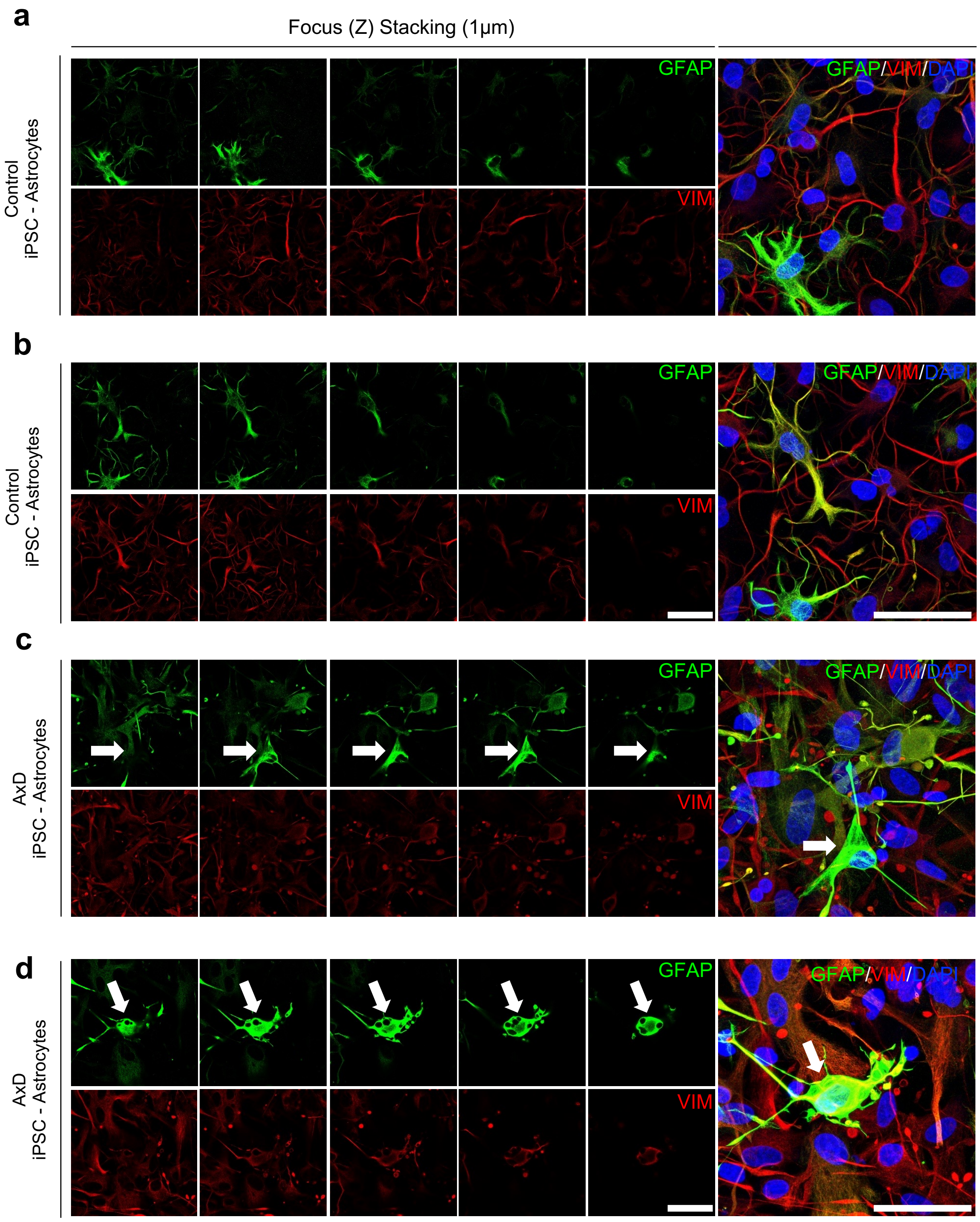
